# Predictive systems biomarkers of response to immune checkpoint inhibitors

**DOI:** 10.1101/2021.02.05.429977

**Authors:** Óscar Lapuente-Santana, Maisa van Genderen, Peter Hilbers, Francesca Finotello, Federica Eduati

**Affiliations:** Department of Biomedical Engineering, Eindhoven University of Technology, Eindhoven, The Netherlands; Biocenter, Institute of Bioinformatics, Medical University of Innsbruck, Innsbruck, Austria; Institute for Complex Molecular Systems, Eindhoven University of Technology, Eindhoven, The Netherlands

## Abstract

Cancer cells can leverage several cell-intrinsic and -extrinsic mechanisms to escape immune system recognition. The inherent complexity of the tumor microenvironment, with its multicellular and dynamic nature, poses great challenges for the extraction of biomarkers of immune response and immunotherapy efficacy. Here, we use RNA-seq data combined with different sources of prior-knowledge to derive system-based signatures of the tumor microenvironment, quantifying immune-cell composition and intra- and inter-cellular communications. We applied multi-task learning to these signatures to predict different hallmarks of immune responses and derive cancer-type-specific models based on interpretable systems biomarkers. By applying our models to independent RNA-seq data from cancer patients treated with PD-1 inhibitors, we demonstrated that our method to Estimate Systems Immune Response (EaSIeR) accurately predicts therapeutic outcome. We anticipate that EaSIeR will be a valuable tool to provide a holistic description of immune responses in complex and dynamic systems such as tumors using available RNA-seq data.

## INTRODUCTION

In the last few years, immunotherapy has revolutionized cancer treatment, especially based on antibodies targeting immune checkpoints such as the cytotoxic T-lymphocyte-associated protein 4 (CTLA-4), programmed cell death protein (PD-1), or its ligand (PD-L1) (Robert, 2020). Immune checkpoint blockers (ICB) boost the patient’s immune system to effectively recognize and attack cancer cells. In different cancer types patients treated with these immune-based therapies have shown promising results, especially in terms of long-term patient survival, and even curative potential. Despite this fact, just a minority of the patients achieve complete response. In addition, high immunological toxicity (Boutros et al., 2016; Postow et al., 2018) and considerable costs (>US$100,000 per patient per year) (Schmidt, 2017) are other challenges for ICB therapy. That is why biomarkers are indispensable to select potential responders and spare unnecessary and potentially harmful treatments to patients that are unlikely to respond to ICB (Tang et al., 2018).

Different mechanisms in the tumor microenvironment (TME) are involved in mediating the immune response and affect the efficacy of ICB therapy (Lapuente-Santana and Eduati, 2020). A first important aspect is the cell type composition present in the TME. Different types of TME cells, and especially immune cells, can have a pro- or anti- tumor role regulating cancer progression and response to treatment (Fridman et al., 2017). A key role in anti-tumor response is played by effector T cells: their state and localization are well known to be essential to assess effectiveness of immunotherapy (Galon and Bruni, 2019). Another important aspect is the inter- and intra- cellular regulation of cellular functions that are responsible for shaping the immune response. Signals from outside the cells are processed by intracellular signalling pathways leading to changes in transcription factors activity and gene expression. Intracellular regulatory networks of tumor cells are involved in innate (endogenous due to mutations) and adaptive (due to exogenous stimulation) mechanisms that tumor cells exploit to resist immune attack (Spranger and Gajewski, 2018). This can be accomplished by different mechanisms such as the upregulation of immune checkpoints (Zerdes et al., 2018), reduced release of inflammatory cytokines (Nagarsheth et al., 2017; Tokunaga et al., 2018) or the impaired antigen presentation by the major histocompatibility complex (MHC) (Cornel et al., 2020). These are all important mechanisms that cancer cells use to communicate with surrounding cells. More in general, ligand-receptor interactions regulate cell-cell communication between all the cells in the microenvironment including tumor cells, immune cells and fibroblasts, and finely regulate tumor characteristics and anti-tumor immune responses (Kumar et al., 2018; Nagarsheth et al., 2017; Ramilowski et al., 2015).

All these aspects should be taken into account to provide a comprehensive description of the TME. A holistic approach to derive biomarkers of immune response can inform clinicians on the efficacy of ICB therapy in individual patients (Lapuente-Santana and Eduati, 2020; Szeto and Finley, 2019). Different emerging *omics* technologies allow to take snapshots of the TME in bulk tumors, single cells from the TME, or from images of tumor tissue slides. The combination of these cutting-edge technologies with new computational tools hold great potential to provide a complete picture of the TME, shedding light on how complex cellular and intercellular mechanisms orchestrate the immune response (Finotello and Eduati, 2018; Finotello et al., 2019a). However, such technologies are not yet widely available and computational tools to fully exploit their potential are still at their infancy. In order to improve precision medicine, we urgently need different approaches to derive a comprehensive description of the TME and how it regulates immune response in individual patients, using currently available patient data. RNA-seq data has become the de facto method to quantify transcriptome-wide gene expression (Stark et al., 2019) and is increasingly available in the clinics (Byron et al., 2016).

Here, we describe an approach based on RNA-seq data combined with different types of prior knowledge to derive a holistic description of the TME. In particular we use computational methods to quantify tumor-infiltrating immune cells (Finotello and Trajanoski, 2018), activity of intracellular signaling and TFs (Garcia-Alonso et al., 2018; Schubert et al., 2018), and extent of intercellular communication (Ramilowski et al., 2015) from bulk-tumor RNA-seq data. Using multi-task machine learning algorithms we aim to assess how these system-based signatures of the TME are associated with 14 different transcriptome-based predictors of anticancer immune responses (**Table S1**), which model different hallmarks of response to ICB therapy.

By training machine learning models on RNA-seq data from 7550 patients’ data across 18 solid cancers generated by The Cancer Genome Atlas (TCGA) (Weinstein et al., 2013), we identified predictive, interpretable system-based biomarkers of immune response in a cancer-type-specific fashion. This integrative approach allowed us to identify several biomarkers that are known to be associated with immune response and response to ICB, as well as new candidates for future follow-up studies. Additionally we showed how the derived system-based biomarkers of immune response are able to predict response to ICB therapy in independent datasets of melanoma (Auslander et al., 2018; Gide et al., 2019) and metastatic gastric (Kim et al., 2018) cancer patients treated with anti-PD-1. This proposed computational framework is provided as a tool called Estimate Systems Immune Response (EaSIeR) that can be applied to bulk-tumor RNA-seq data to investigate mechanistic biomarkers and predict patients’ likelihood to respond to ICBs.

## RESULTS

### Multiple views of the tumor microenvironment

Using bulk RNA-seq data combined with different types of biological prior knowledge, we derived five types of system-based signatures of the TME for 7550 cancer patients across 18 solid cancers from the TCGA data as summarized in **Figure 1A** (additional details in **STAR Methods**).

**Figure 1:**
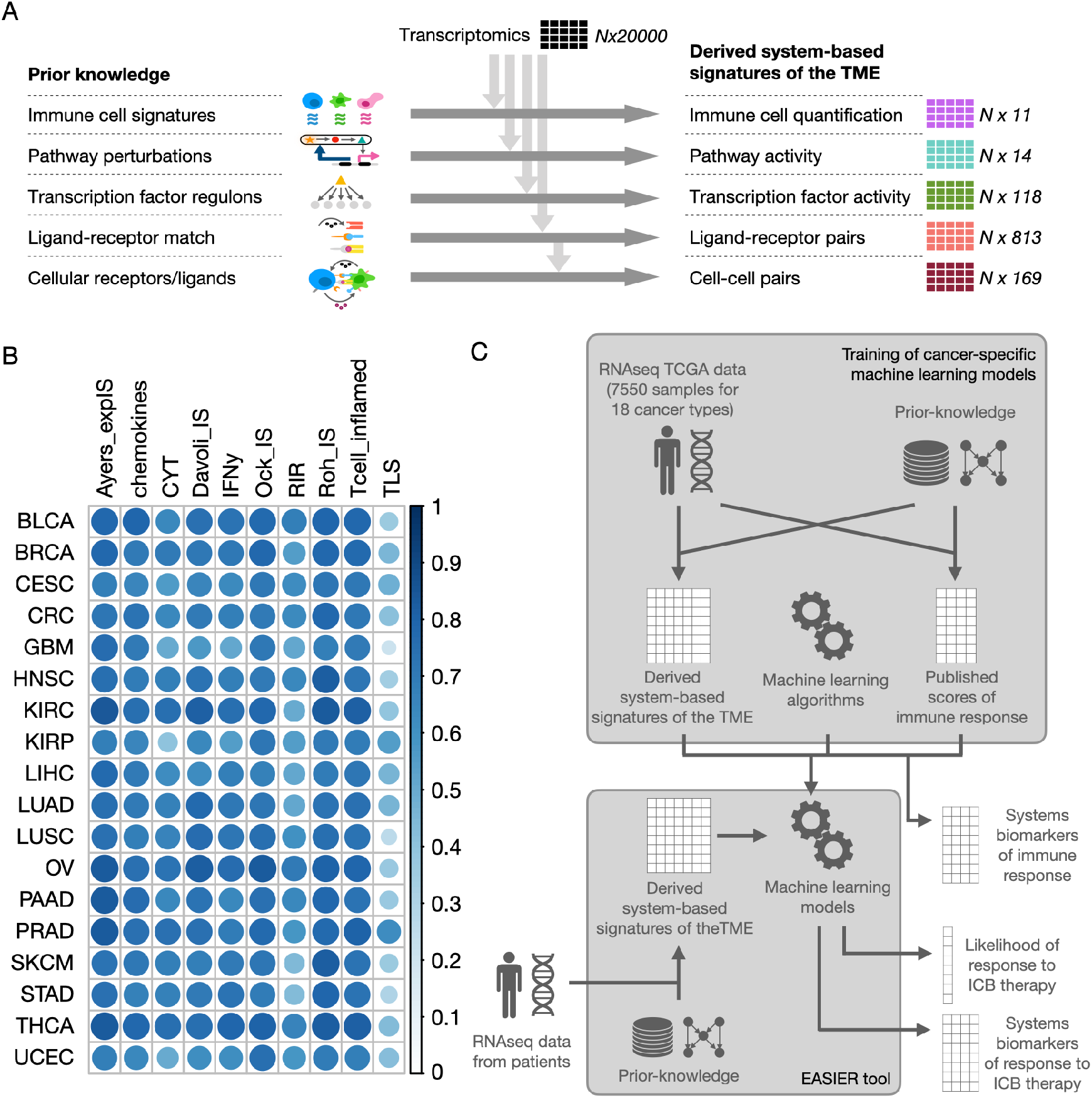
Overall description of the approach taken. (A) Derivation of the five system-based signatures of the TME based on the integration of RNA-seq data and different sources of prior knowledge. (B) Cancer-specific median correlation of each of the 10 used scores (described in **Table S1**) of immune response with all the other 14 scores. (C) Schematic pipelines. Cancer-specific models are trained on TCGA data. System-based signatures of the TME and scores of immune response are derived combining RNA-seq data and prior knowledge, and are used respectively as algorithm inputs and output. Trained models are used to define systems biomarkers of immune response. These models are also included in a tool called EaSIeR that allows users to input RNA-seq data for new patients and to compute the likelihood of response to immunotherapy and the biomarkers distinguishing responders and non-responders.

The first type of signature consists of immune cell fractions obtained with quanTIseq (Finotello et al., 2019b). quanTIseq cell fractions are estimated using a deconvolution approach leveraging as prior knowledge cell-type specific expression signatures for: B cells, classically (M1) and alternatively (M2) activated macrophages, monocytes (Mono), neutrophils (Neu), natural killer (NK) cells, non-regulatory CD4^+^ T cells, CD8^+^ T cells, regulatory T (Treg) cells, and myeloid dendritic cells (DCs). quanTIseq also provides the fraction of ‘other’ unclassified cells in the mixture, resulting in a total of 11 cellular features.

We considered two types of signatures describing intracellular networks, quantifying the activity of 14 signaling pathways and 118 transcription factors (TFs). Pathway activity was derived using PROGENy (Holland et al., 2020; Schubert et al., 2018), which uses as prior knowledge perturbation-response gene signatures extracted measuring downstream gene changes upon perturbations of a pathway. The activity of the pathways is computed as linear combinations of their signature genes. TFs activity was computed using DoRothEA (Garcia-Alonso et al., 2018), which assumes as prior knowledge the networks of TF-targets interactions (regulons) and infers the activity of each TF from the expression of its target genes.

Additionally, we extracted two types of signatures related to inter-cellular networks, quantifying 813 ligand-receptor (LR) pairs and 169 cell-cell (CC) pairs. To compute LR pairs we leveraged as prior-knowledge 1894 LR pairs annotated in (Ramilowski et al., 2015). We filtered for literature-supported pairs expressed in 25 cell-types that are present in the TME, including immune cells, cancer cells, fibroblasts, endothelial cells and adipocytes (**STAR Methods**). The weight for the LR pairs was computed as the limiting factor between the expression of the ligand and the receptor. We then computed a score of the cell-cell interactions between 13 aggregated cell types (including autocrine signaling), as a weighted sum of the number of LR pairs expressed for each CC pair (**STAR Methods**).

For the same TCGA samples we also computed 14 different transcriptomics-based scores of immune response (**Table S1**). All these scores were recently published and have been proposed as predictors of response to ICB therapy. We computed cancer-specific correlations between the 14 scores (**Figure S1**) and identified a subset of 10 scores that were highly correlated across all 18 cancer types (across cancer types median of the median Pearson correlation with all other scores > 0.4, **Figure 1B**).

We considered these scores as output variables (*tasks*) and trained two different multi-task machine learning algorithms using the derived system-based signatures as input features (**Figure 1C**). Multi-task learning allows solving multiple learning tasks at the same time, exploiting the shared information between tasks. Therefore, only the 10 correlated scores of immune response were used as tasks for model training. The first approach that we used is regularized multi-task linear regression (RMTLR) (Yuan and Lin, 2006a) using Elastic Net regularization (**STAR Methods**). Regularization allows to improve model generalization (avoiding overfitting on the training data) and to perform selection of relevant predictive features (common for all tasks) which we interpreted as systems biomarkers of immune response. The second approach that we used is Bayesian Efficient Multiple Kernel Learning (BEMKL) (Gönen, 2012), which was the best performing algorithm in the NCI/DREAM7 challenge on prediction of cell lines drug sensitivity from genomic information (Costello et al., 2014). While BEMKL is a more sophisticated approach that can account for non-linearities, it does not allow us to directly select the important predictive features. Cancer-specific models were trained using RMTLR and BEMKL with randomized cross-validation using as input data each of the five system-based signatures separately, pairwise combinations and combination of all views (cross-validation performances in **Figure S2**). For RMTLR, the randomized cross-validation was also used to select only robust features (**STAR Methods**). The trained cancer-specific models are provided as a tool called EaSIeR (**STAR Methods**). Users can provide RNA-seq data and use the tool to derive system-based signatures of the TME, analyze systems biomarkers of immune response and predict patient-specific likelihood of response to ICB therapy (**Figure 1C**).

For models optimized using RMTLR, we computed the median across cancer-types of the estimated feature weights and verified that 99% of the feature weights (estimated separately for each view) had variance ≤ 0.0015 across tasks, proving that biomarkers are in general consistent across tasks (**Figure S3**). By clustering tasks based on feature weights, we obtained four main clusters: 1) cytolytic activity (CYT) (Rooney et al., 2015); 2) Tertiary Lymphoid Structures (TLS) signature (Cabrita et al., 2020); 3) chemokines (Messina et al., 2012) and IFNy (Ayers et al., 2017) signatures (both related to cytokine production); and 4) all remaining six immune signatures (**Figure S3**).

We analyzed the systems biomarkers that we identified using RMTLR separately for each type of system-based signatures of the TME, i.e. immune cells, intracellular networks (pathways and TFs activity) and intercellular networks (LR and CC pairs). Then we assessed the performance of our models in predicting response to ICB therapy on independent datasets, analyzed systems biomarkers that differentiate responders and non-responders to therapy, and evaluated the effect of combining different types of signatures. Finally, we performed a preliminary analysis on the potential of integrating into the models orthogonal information from imaging and proteomics data. The results of these analyses are presented in the following sections.

### Immune cells as biomarkers of immune response

We identified several relevant robust associations between immune cell composition and scores of immune response (**Figure 2**; **STAR Methods**). In particular, CD8^+^ T cells, which are essential for tumor-cell recognition and killing (Chen and Mellman, 2013), were identified as positive biomarkers for all cancer types (**Figure 2A**). The fraction of ‘other’ uncharacterized cells positively correlates with tumor purity and negatively correlates with the percentage of tumor-infiltrating infiltrating immune cells (Finotello et al., 2019b). Here, we observed a negative correlation of this feature with immune response across all cancers, which can be interpreted as a positive association between immune infiltration levels and immune response (**Figure 2A**). Some immune cells such as T_reg_ cells, M1 and M2 macrophages, and B cells were positively associated with response in most cancer types (16, 17, 16 and 14 respectively, out of the 18 analyzed cancer types; **Figure 2A**). For most of the cell types the association was consistent across all four clusters of tasks (**Figure 2B**), with the exception of B cells, CD8^+^ T cells and M1 macrophages. B cells showed a particularly strong association with the TLS signature. TLS are organized lymphoid aggregates and recent studies suggested that B cells are localized in TLS, and that B cells and TLS contribute to an effective T cell response to ICB (Cabrita et al., 2020; Helmink et al., 2020). Instead, CD8^+^ T cells and M1 macrophages are respectively less strongly or mildly negatively associated with TLS.

**Figure 2:**
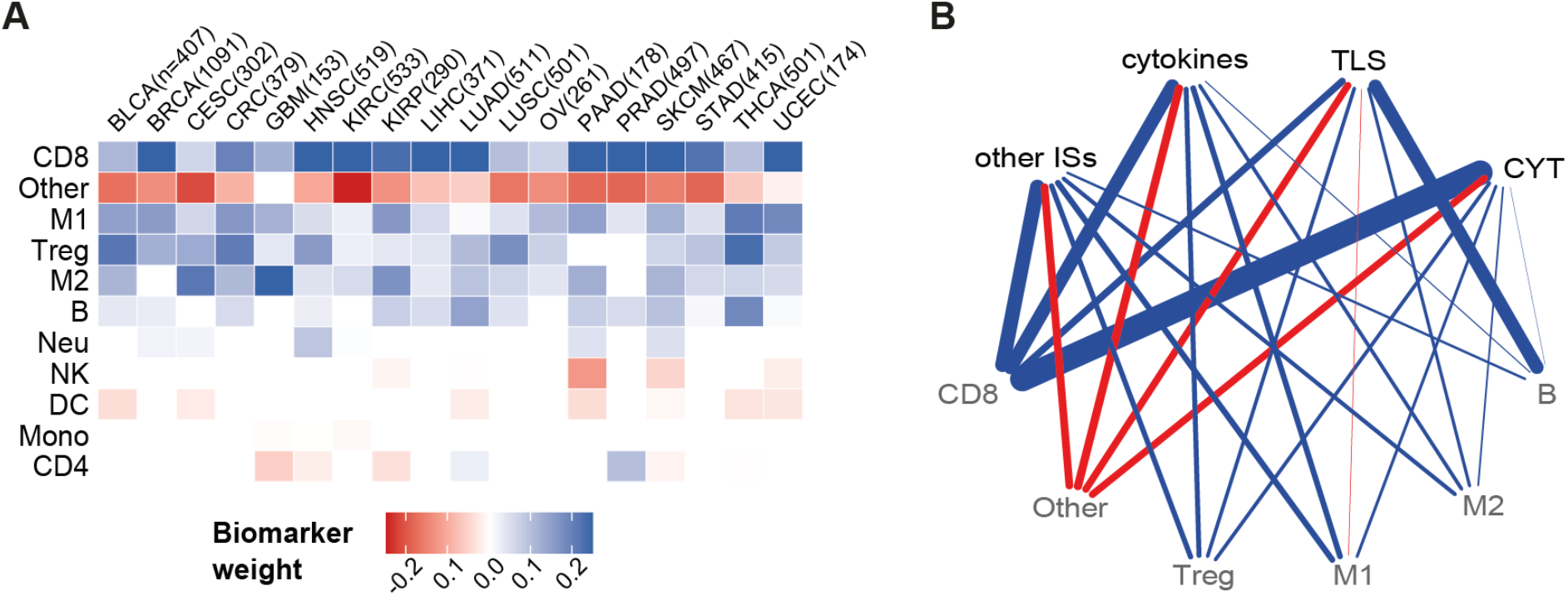
Systems biomarkers based on immune cells quantification. (A) Heatmap showing regression coefficients for cancer type-specific models when using immune cells quantification as biomarkers. Shown are the median values computed first across 100 randomized cross-validation runs (to keep only robust biomarkers) and then across tasks. Rows (biomarkers) were sorted according to their absolute mean value across tumors. (B) Network representing associations between clusters of tasks (top nodes) and immune cell biomarkers (bottom nodes). Only top five biomarkers for each cluster of tasks (ranked by median weight across the tasks in the cluster) are shown. Edge widths represent the median weight of each biomarker across cancer types. Positive (blue), none (white) or negative (red) association of each biomarker with the tasks which are hallmarks of immune response. B, B cells; CD4, non-regulatory CD4^+^ T cells; CD8, CD8^+^ T cells; DC, dendritic cells; M1, classically activated macrophages; M2, alternatively activated macrophages; Mono, monocytes; Neu, neutrophils; NK, natural killer cells; Treg, regulatory T cells.

### Intracellular networks as biomarkers of immune response

Tumor-cell intrinsic deregulation of cellular signaling due to mutations or epigenetic alterations have an effect on the functioning of intracellular networks that regulate cellular phenotype but also on the interaction with the immune system (Spranger and Gajewski, 2018). Among the analyzed pathways and TFs activities we identified several biomarkers of immune response (**Figure 3**).

**Figure 3:**
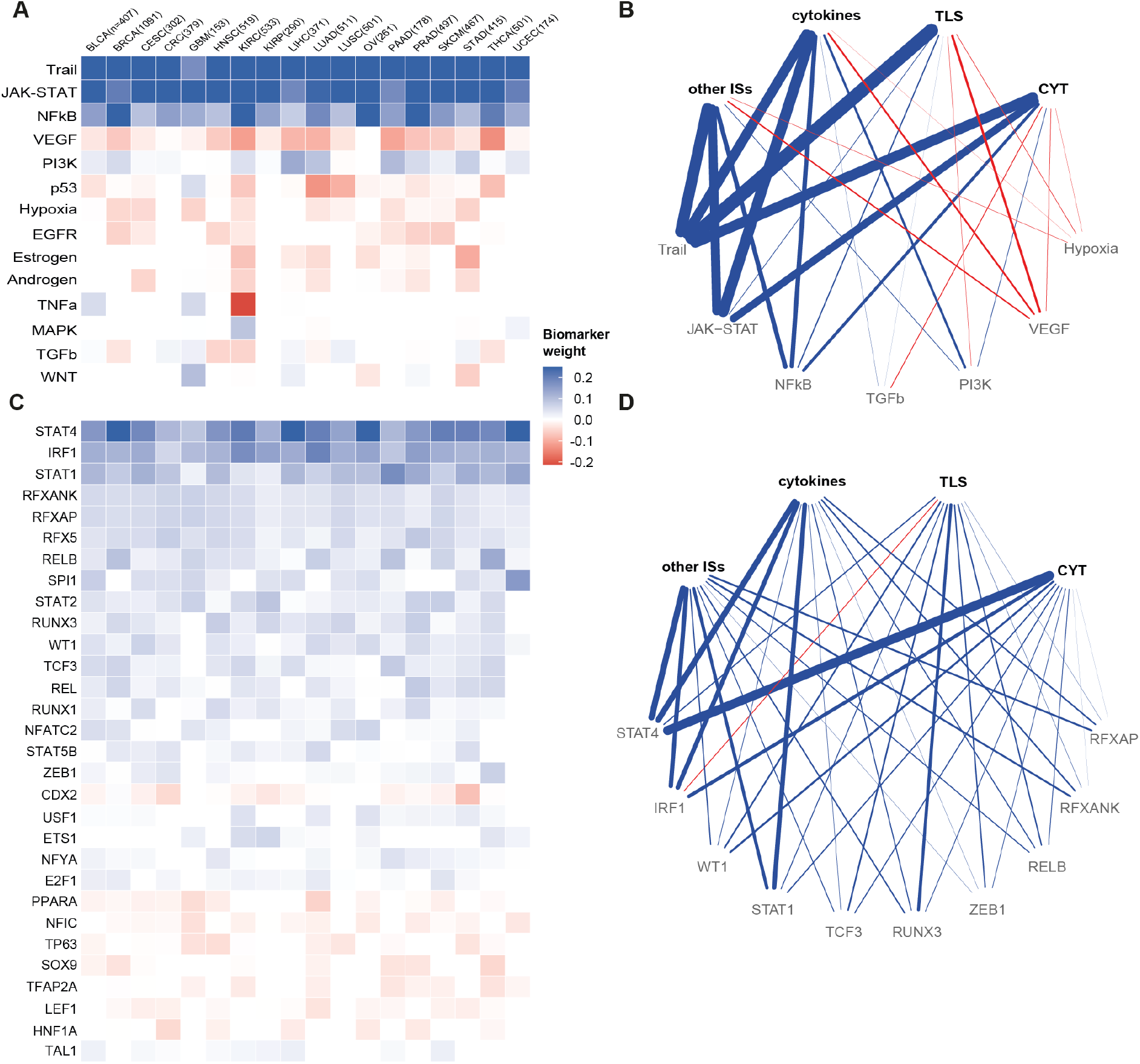
Systems biomarkers based on pathway and TF activity. Heatmap showing regression coefficients for cancer type-specific models when using (A) pathways activity and (C) TF activity (limited to the top 30) as biomarkers. Shown are the median values computed first across 100 randomized cross-validation runs (to keep only robust biomarkers) and then across tasks. Rows (biomarkers) were sorted according to their absolute mean value across tumors. Network representing associations between clusters of tasks (top nodes) and biomarkers (bottom nodes) from (B) pathway and (D) TF activity. Only top five biomarkers for each cluster of tasks (ranked by median weight across the tasks in the cluster) are shown. Edge widths represent the median weight of each biomarker across cancer types. Positive (blue), none (white) or negative (red) relationship of each biomarker with the tasks which are hallmarks of immune response.

We identified a strong positive association between TRAIL apoptotic pathway, JAK-STAT, and NFkB pathways and the predicted immune responses in all cancer types (**Figure 3A**; **STAR Methods**). The TRAIL pathway can be activated by different types of immune cells causing tumor cell apoptosis (Johnstone et al., 2008). In turn, tumor cell death results in the activation of the immune system via the cancer-immunity cycle (Chen and Mellman, 2013). The JAK-STAT and NFkB pathways are also known to play pivotal roles in immune responses. Both pathways are stimulated by IFNγ released mainly by NK and T cells (Jorgovanovic et al., 2020) and regulate several mechanisms of adaptive immune resistance, including upregulation of immune checkpoint molecules (Zerdes et al., 2018), inhibition of productions of pro-inflammatory chemokines (Ding et al., 2016), and promoting expression of class-I MHC molecule expression (Cornel et al., 2020). Both JAK-STAT and NFkB pathways showed a lower association with TLS than with other tasks (**Figure 3B**), suggesting that their role in immune response is not dependent on TLS.

Activation of the PI3K pathway, which we identified as a biomarker for 16 cancer types (**Figure 3A**), was also shown to enhance PD-L1 expression (Parsa et al., 2007; Vivanco and Sawyers, 2002). PI3K pathway activation can be caused by different mechanisms of innate resistance to immune response, such as loss of PTEN, which is an inhibitor of PI3K pathway, or oncogenic mutation of PIK3CA gene. Direct or indirect therapeutic inhibition of PI3K was shown to reduce PD-L1 expression and increase anti-tumor immunity (Zerdes et al., 2018).

The positive biomarkers described above, are pathways generally associated with inflamed tumors, which are usually more predisposed to respond to ICB immunotherapy (Chen and Mellman, 2017). In contrast, for VEGF we observed a negative association with immune response for 17 cancer types, and no association for ovarian cancer (OV) (**Figure 3A**). The negative association is in line with the role that VEGF plays in promoting immune-exclusion due to the presence of vascular barriers (Chen and Mellman, 2017). Immune-excluded tumors are less responsive to ICB therapy (Bonaventura et al., 2019), and inhibition of VEGF could promote immune infiltration and improve efficacy when used in combination with ICB therapy (Manegold et al., 2017).

Similarly, the p53, hypoxia, and EGFR pathways also showed negative correlation for the majority of the cancer types (14, 13 and 9 respectively, **Figure 3A**). Interestingly, it has been recently shown that, in lung cancer, loss-of-function mutations in the tumor suppressor P53 gene is associated with increased expression of PD-L1, immune cell infiltration, and tumor immunogenicity and may help predicting response to ICB therapy (Dong et al., 2017). Our findings suggest that the activity of the p53-mediated DNA damage response pathway might be considered as a predictor of ICB therapy response as well, not only for lung cancer (LUAD and LUSC have the strongest associations) but also for other cancer types (**Figure 3A**). Notably, p53 pathway revealed a positive correlation in GBM, this is in agreement with stronger immune responses found in TP53 wild-type GBM patients compared to patients harboring TP53 mutations (Qin et al., 2020).

We also observed a negative impact of hypoxia on the immune response (**Figure 3A**), consistent with recent observations that hypoxia impairs the anti-tumor immunity and contributes to resistance to immunotherapy (Noman et al., 2019). Preliminary studies in mice revealed the potential of targeting hypoxia in combination with ICB therapy to restore T cell infiltration and increase efficacy of immunotherapy (Jayaprakash et al., 2018).

Consistently with our results (**Figure 3A**), it has been shown that activation of the EGFR signaling pathway contributes to an uninflamed tumor microenvironment, and that combining EGFR inhibitors and anti-PD-1/PD-L1 antibodies could improve therapeutic outcome in EGFR-mutant tumors (Yu et al., 2018).

Next, we focused our analysis on the association between TF activity and immune responses (**Figure 3C** for the top 30 biomarkers, full list in **Table S2**). We identified several TFs which were selected consistently across the majority of cancer types. For instance, STAT1, STAT2, and STAT4, all selected as positive biomarkers (**Figure 3C**), are members of the STAT family in the JAK-STAT signaling pathway discussed above (Garcia-Diaz et al., 2019). Although STAT3 is often considered an important player in cancer immunotherapy (Zou et al., 2020), it was not selected as top biomarker in our analysis. This is in line with recent publications suggesting that the main role in regulation of PD-L1 expression is played by STAT1 (Bellucci et al., 2015). In tumor cell lines from several cancer types, siRNA knockdown of STAT3 did not reduce IFNγ and IL27 induced PD-L1 protein expression, while siRNA knockdown of STAT1 did (Chen et al., 2019). STAT4 deficiency has been associated with diminished anti-tumor immune response and worse prognosis (Anderson et al., 2019; Nishi et al., 2017). The positive association of STAT4 found in all the 18 cancer types we analyzed suggests that this mechanism might play a major role in anti-cancer immunity and, possibly, response to ICB pan-cancer. As previously observed for the JAK-STAT pathway, also the STAT TFs seem to be not dependent on TLS (**Figure 3D**).

Another relevant biomarker (selected for all cancer types, **Figure 3C**) is IRF1. The IRF1 TF can regulate the expression of PD-L1 (Garcia-Diaz et al., 2019; Lee et al., 2006) and the productions of cytokines, including the CXCL9 chemokine that is responsible for recruiting anti-tumor immune cells (Ding et al., 2016). Similarly, also RELB (selected for 17 out of 18 cancer types), which is part of the NFkB complex, regulates PD-L1 expression and inflammation (Gowrishankar et al., 2015). RELB also regulates MHC-I gene transcription (Cornel et al., 2020). Tumor-immune infiltration favored by pro-inflammatory cytokines, susceptibility of cancer cells to immune-effector mechanisms such as MHC-I genes expression, and expression of PD-L1 are all hallmarks of effective immunotherapy (Galluzzi et al., 2018).

Other important positive biomarkers shared across all 18 cancer types were RFXANK, RFXAP, and RFX5, which form the RFX trimeric complex (**Figure 3C**). This complex cooperates with NLRC5 to drive the transcription of class-I MHC genes (Meissner et al., 2012). Accordingly, recent studies suggested that reduced activity of NLRC5 plays a key role in immune evasion (Chelbi and Guarda, 2016; Yoshihama et al., 2017). Taken together, these results may hint at RFX as a candidate biomarker of anti-tumor immunity.

Another positive biomarker was RUNX3 (selected for 16 cancer types, **Figure 3C**), which plays a role in the tumor microenvironment regulating hypoxia and immune-cell infiltration, and has been suggested as a potential target to prevent immune escape of cancer cells (Manandhar and Lee, 2018). We observed that RUNX3 was more strongly associated with TLS than with other immune signatures (**Figure 3D**).

Additionally, we found a number of biomarkers negatively associated with immune response, although the weight of the association was in general lower. CDX2 was identified as a negative biomarker in 14 cancer types (**Figure 3C**). Loss of CDX2 has been reported in colorectal (CRC) tumors that are microsatellite unstable (Aasebø et al., 2020) or PD-L1 positive (Inaguma et al., 2017). These observations are in agreement with the negative association that we identified and suggest a potential role of CDX2 as a biomarker of ICB therapy. Other results suggested that CDX2 might play an important role also in other cancer types and in particular in stomach cancer (STAD). In our results, CRC and STAD showed the strongest negative associations with immune response for CDX2, but a negative association was identified also for other cancer types (**Figure 3C**). Another interesting example of negative biomarker (for 11 out of the 18 cancer types, **Figure 3C**) was PPARA, which encodes for a ligand-activated TF regulating lipid metabolism and fatty acid oxidation (Bougarne et al., 2018). The PPARA antagonist TPST-1120 promotes a more inflamed tumor microenvironment in different types of cancer and is currently in clinical trial as monotherapy and in combination with anti-PD1 therapy (Laport et al., 2019). Blocking PPARA shifts the metabolic balance of immune cells from fatty acid oxidation towards glycolysis, which works in favor of certain immune cells cell populations such as M1 macrophages and effector T cells but against M2 macrophages and T_reg_ cells (Laport et al., 2019). This was in line with the negative association that we observed for PPARA, because an inflamed microenvironment is essential for effective ICB therapy.

### Biomarkers of immune response based on cell-cell communication

The phenotype of cancer cells is not only defined by intracellular oncogenic pathways but also by their exchange of signals with other TME cells. We, therefore, analyzed the potential of intercellular data in driving an effective immunotherapy response through ligand-receptor (LR) and cell-cell (CC) interactions among cell types of the TME (**Figure 4**; **STAR Methods**).

**Figure 4:**
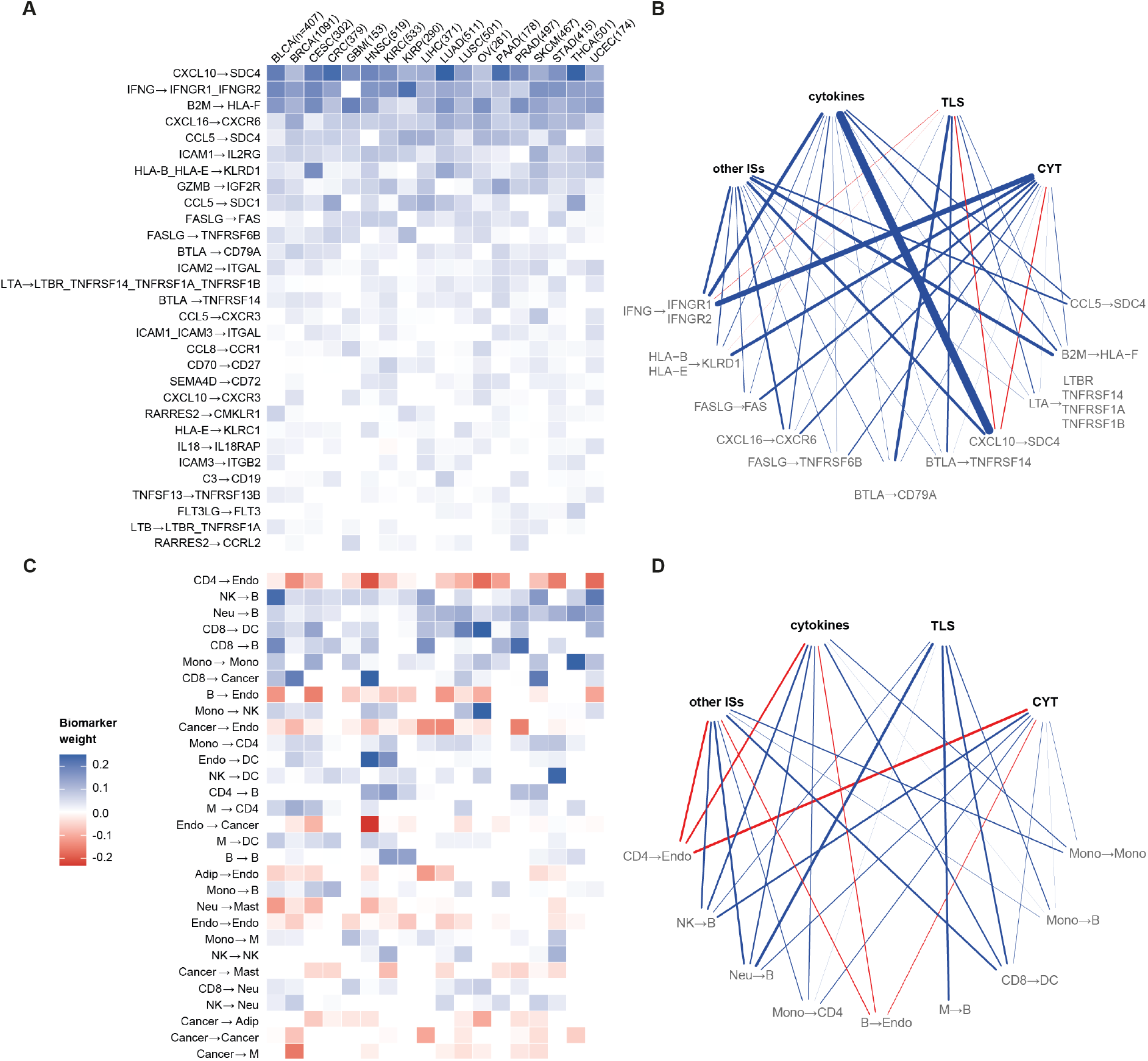
Systems biomarkers based on cell-cell interactions. Heatmap showing top 30 regression coefficients for cancer type-specific models when using (A) ligand-receptors pairs and (C) cell-cell pairs as biomarkers. Shown are the median values computed first across 100 randomized cross-validation runs (to keep only robust biomarkers) and then across tasks. Rows (biomarkers) were sorted according to their absolute mean value across tumors. Network representing associations between clusters of tasks (top nodes) and biomarkers (bottom nodes) from (B) ligand-receptor and (D) cell-cell pairs. Only top five biomarkers for each cluster of tasks (ranked by median weight across the tasks in the cluster) are shown. Edge widths represent the median weight of each biomarker across cancer types. Positive (blue), none (white) or negative (red) relationship of each biomarker with the tasks which are hallmarks of immune response. Arrows in biomarkers names indicate the direction of the interaction (including cases of autocrine signaling).

Among the LR (sender→receiver molecule) biomarkers (top 30 biomarkers in **Figure 4A**, full list in **Table S3**), we found several pairs of chemokines and corresponding receptors that are important for attracting immune cells to the TME (Nagarsheth et al., 2017). An important chemokine in the TME is CXCL10, which regulates immune cell migration, differentiation, and activation (Tokunaga et al., 2018). Higher levels of CXCL10 are associated with increased number of infiltrated CD8^+^ T cells (Nagarsheth et al., 2017). The most studied receptor of CXCL10 is CXCR3 (LR pair associated with immune response in 12 out of the 18 cancer types, **Figure 4A**), which is expressed by effector CD8^+^ T cells, T helper 1 (Th1) cells and NK cells, which are all anti-tumor lymphocytes (Nagarsheth et al., 2017). However, the top CXCL10-related LR pair, which we identified as a positive biomarker for all cancer types, is CXCL10→SDC4 (**Figure 4A**). While there is limited evidence on the role of SDC4 in cancer, it has been shown that interaction between CXCL10 and SDC4 inhibits fibroblast recruitment in pulmonary fibrosis (Jiang et al., 2010). Our results suggest that a similar mechanism might take place in the tumor, since fibroblasts recruited in the TME can suppress immune response and limit ICB immunotherapy efficacy (Barrett and Puré, 2020). In agreement with the role of CXCL10, the CXCL10→SDC4 pair was particularly strongly positively associated with cytokines related immune signatures, while showing a small negative association with TLS and CYT (**Figure 4B**).

CCL5 is another chemokine that we identified as positively associated with immune response when bound to different receptors (SDC4, SDC1, and CXCR3 for 16, 15, and 13 cancer types respectively, **Figure 4A**). CCL5 binding to CCR5 is often described as a main actor of tumor progression (Aldinucci et al., 2020). However, recent studies have also shown that CCL5 overexpression can favor CD8^+^ T cell infiltration, anti-tumor immunity, and immunotherapy response (Dangaj et al., 2019) in agreement with the positive association that we identified. Similarly, we found the CXCL16→CXCR6 LR pair as positively associated with immune response for all 18 cancer types. Although the role of CXCL16 in cancer is disputed (Kim et al., 2019), overexpression of CXCL16 by tumor cells is associated with increased infiltration of CD8^+^ T cells (Hojo et al., 2007) and NK cells (Yoon et al., 2016).

Another relevant LR biomarker was IFNG binding to IFNGR1 and IFNGR2 (positive association for 17 of the 18 cancer types, **Figure 4A**). IFNG is the gene encoding for the IFNγ cytokine that, as already discussed in the pathways section, plays a main role in immune response (Sharma et al., 2017). Mutations in the interferon gamma pathway (including IFNGR1 and IFNGR2) are associated with resistance to ICB therapy (Gao et al., 2016). Also in this case, we observed that IFNγ-dependent mechanisms of immune response are independent of TLS (**Figure 4B**).

We also observed several positive biomarkers related to T-cell-mediated cancer cell killing (**Figure 4A**). These include the GZMB gene, which encode the granzyme B serine protease that is secreted by CD8^+^ T cells and NK cells and induces apoptosis in target cells binding to the corresponding receptor IGF2R (Motyka et al., 2000). In line with our results, upregulation of the receptor on the membrane of tumor cells promotes penetration of granzyme B favoring immune cell-mediated apoptosis and was suggested as a potential target for immunotherapy (Li et al., 2017). Similarly, Fas ligand (encoded by FASLG gene) binding to the TNFRSF6/FAS receptor (part of the TNF receptor superfamily) is involved in CD8^+^ T cells and NK cell-mediated apoptosis (Modiano and Bellgrau, 2016). Another interesting apoptosis-related biomarker is BTLA, which is a ligand for CD79A and TNFRSF14. Binding of BTLA to TNFRSF14 (also known as HVEM) is an immune checkpoint that has been generally associated with negative immune responses (Serriari et al., 2010), albeit there is also evidence that this binding promotes survival of CD8^+^ T cells in melanoma (Haymaker et al., 2015). Another member of the TNFR superfamily that we identified as a biomarker is CD27, which expressed by CD8^+^ T cells binds to its receptor CD70 on antigen presenting cells upon T-cell activation. The CD27/CD70 axis can be targeted with antibodies and its blockade has potential for cancer immunotherapy (van de Ven and Borst, 2015).

We additionally identified as positive biomarkers several interactions between MHC-I genes and the corresponding receptors (HLA-B_HLA-E→KLRD1 and HLA-E→KLRC1 in 17 and 14 cancer types respectively). These receptor genes encode for CD94/NKG2, which is a family of inhibitory receptors expressed mainly on the surface of NK and CD8^+^ T cells (André et al., 2018). Anti-NKG2A antibody is a checkpoint inhibitor in clinical trials that was reported to enhance tumor immunity by promoting functioning of these immune cells (André et al., 2018). Another relevant positive biomarker for all 18 cancer types is B2M→HLA-F. Although this is an intracellular interaction rather than a LR pair, the importance of this interaction for immune response is clear since B2M stabilizes the MHC-I complex allowing recognition by T-cell receptor (Del Campo et al., 2014).

Another important mechanism that we identified among the LR biomarkers, is the stimulation of LFA-1 (encoded by ITGAL and ITGB2 genes) by ICAM (ICAM2→ITGAL, ICAM1_ICAM3→ITGAL and ICAM3→ITGB2 in 18, 11 and 8 cancer types respectively). LFA-1 is essential for adhesion of CD8+ T cells and NK cells to the cancer cell, therefore allowing their activation (Reina and Espel, 2017). On mouse models, the ICAM1-LFA-1 interaction was also shown to cause clusters of activated T-cells in the tumor and was suggested as a mechanism of tumor mediated immune retention that prevents trafficking of T cells to lymph nodes (Yanguas et al., 2018). In the same paper they suggest ICAM-1 as a potential target for cancer treatment by incrementing lymphocyte migration to the lymph nodes.

We went further and analyzed the number of active LR interactions per cell-cell (CC) pair, generating a score for each CC (sender→receiver cell) pair. CC scores were used to build a model that allowed us to disentangle the complex crosstalk between cells of the TME and its influence on immune responses (top 30 biomarkers in **Figure 4C**, full list in **Table S4**).

CC communication profiles were very specific for each cancer type, with no biomarker shared across all cancer types (**Figure 4C**). The sign of the association with immune response however tended to be consistent across the different cancer types. Endothelial cells appeared as receiver cells for 5 of the top 30 CC pairs biomarkers (with CD4^+^ T cells, B cells, cancer cells, adipocytes, and endothelial cells as sender cells). In all cases they had a consistent, negative association for all cancer types for which they were selected as biomarkers. The weight of this association was consistent for all tasks except TLS that showed no association (**Figure 4D**). Endothelial cells contributed to establishing an immunosuppressive TME, being actively involved in immune cell exclusion and inhibition of lymphocyte activation (Klein, 2018). The negative association that we identified is in line with the fact that inhibition of endothelial cells favour an anti-tumor immune response (Klein, 2018). As expected, signalling of CD8^+^ T cells to dendritic cells and cancer cells was identified as strong positive biomarker (in 14 and 10 of the 18 cancer types respectively, **Figure 4C**), in line with the crucial role these cells play in immune response and mediation of immunotherapy effects (Raskov et al., 2021). Among the top 30 biomarkers, we also found B cells as receiver cells in 6 different CC pairs (with NK cells, Neutrophils, CD8^+^ T cells, CD4^+^ T cells, B cells as sender cells and monocytes). In all cases, these CC pairs were positively associated with immune response, which is in line with the role of B cells as popular factories of antibodies after antigen recognition. In particular, NK cells and Neutrophils help B cells by regulating their activation (NKcells→B-Cell and Neutrophils→B-Cell were selected in 15 and 16 cancer types; **Figure 4C**) (Blanca et al., 2001; Parsa et al., 2016).

### Prediction of response to immunotherapy with PD-1 blockers

We next assessed the performance of EaSIeR on independent RNA-seq data from patients with melanoma (Auslander et al., 2018; Gide et al., 2019) and metastatic gastric cancer (Kim et al., 2018) treated with anti-PD1 immunotherapy (**STAR Methods**; **Table S5**). Pre-therapy RNA-seq data were provided as input to EaSIeR, which derived patent-specific, system-based signatures of the TME. The cancer-type-specific machine learning models built on the TCGA data (using BEMKL and RMTLR methods, **STAR Methods**) were then used to predict patient likelihood to respond to ICB therapy.

First, we assessed model performances in stratifying patients into responders (R) and non-responders (NR) (**Figure 5A-F**). For this, we used models built separately on each of the five described system-based signatures of the TME (single views), pairwise combinations of views, and combination of all views.

**Figure 5:**
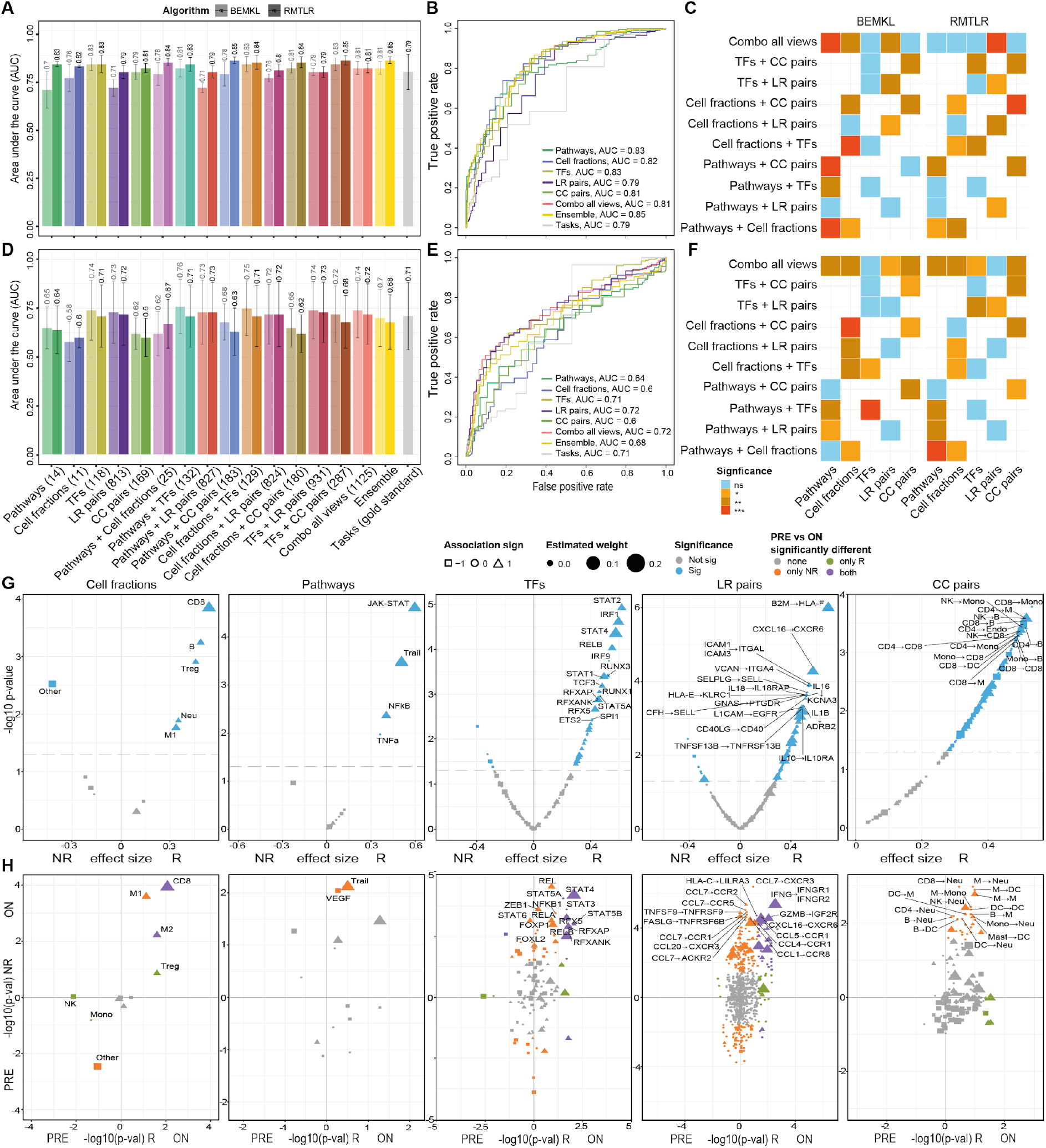
EaSIeR evaluation on independent datasets. Area Under the Curve (AUC) values for the melanoma datasets (Auslander et al., 2018; Gide et al., 2019) (A) and for the metastatic gastric cancer dataset (Kim et al., 2018) (D) based on system-based signatures of the TME considering: single views, pairwise combinations of views, combination of all views, average of single views predictions (ensemble), and average of all tasks (gold standard). Barplots represent the average AUC across tasks and error bars describe the corresponding standard deviation. Receiver operating characteristic (ROC) curves for the melanoma (B) metastatic gastric cancer (E) datasets based on system-based signatures of the TME (single views), combination of all views, average of single views predictions (ensemble), and average of all tasks (gold standard). ROC curves and AUC values are computed using the mean of the predictions across tasks. Performance comparison between single (x axis) and combined (y axis) views for the melanoma (C) metastatic gastric cancer datasets, using BEMKL (left) and RMTLR (right) algorithms (one-side Wilcoxon signed-rank test). Statistical significance is indicated by colors according to the legend. The significant level (*p-value < 0.05, **p-value < 0.01, ***p-value < 0.001) indicates whether combining views improves the performance. (G) Volcano plots for systems biomarkers of the immune response from the melanoma datasets comparing non-responder (NR) and responder (R) patients (two-sided Wilcoxon-rank sum test). Significant biomarkers (p-value < 0.05) are shown in blue. Biomarkers are drawn according to their corresponding sign (shape) and weight (size) obtained during model training. Labels are reported for the top 15 significant biomarkers. (H) Starburst plots showing the statistical comparison (−log_10_ p-value, two-sided Wilcoxon-rank sum test) between pre- and on-treatment samples for responders (R, x-axis) and non-responders (NR, y-axis). The sign is used to show if the biomarkers are higher ON (positive sign) or PRE (negative sign) treatment. Biomarkers are colored according to their consistent statistical significance in both NR and R patients. Labels are reported for the top 15 significant biomarkers.

On the melanoma datasets, RMTLR applied to the single views was able to accurately predict patient response (average AUC = [0.79 - 0.84]), with performance comparable or superior to the gold standard computed as the average of all the tasks (average AUC = 0.79; **Figure 5A-B**). In particular, the ensemble model (average AUC = 0.85), computed as the average of the predictions from the single views, performed significantly better than the average of the literature-based tasks (p-value = 0.003, effect size = 0.849). Performances using BEMKL were in general inferior. This might be due to the fact that BEMKL is designed to handle problems with a large number of features (**STAR Methods**), while all the single views considered here had a relatively low number of features (from few dozens to hundreds, compared to thousands used in previous applications of the BEMKL model (Ali et al., 2018; Costello et al., 2014)). Combining pairs of different views significantly improved performances, in particular for: cell fractions + CC pairs, pathways + cell fractions, pathways + CC pairs and cell fractions + TF for RMTLR (**Figure 5C**; **STAR Methods**). Particularly good predictions were obtained combining information on pathways and cell fractions (average AUC using RMTLR = 0.84), despite the very limited number of features used (25 in total), performing even better than the combination of all views (average AUC using RMTLR = 0.81). Performances on the metastatic gastric cancer dataset were in general inferior compared to the results obtained on the melanoma dataset (single views average AUC range 0.6-0.74 and 0.58-0.74 using RMTLR and BEMKL respectively), in line with the lower performances of the gold standard (average AUC = 0.71, **Figure 5D**). Reduced performances might be due to the fact that models were trained on non-metastatic samples.

Unlike previous predictors that are based on simple gene sets, EaSIeR systems-biology approach allows investigating the mechanisms behind the differential patients’ response to treatment. For the melanoma dataset, we investigated how the identified systems biomarkers differ between responding and non-responding patients and evolve upon treatment (comparing pre- and on-treatment data). As expected, the top biomarkers for all individual views were the best at discriminating between responders and non-responders (**Figure 5G**). Responders had a higher number of infiltrated immune cells, including CD8^+^ T cells, and a lower number of ‘other’ non-immune cells. They also had higher activity in pathways (e.g. JAK-STAT, NFkB) and TFs (e.g. STAT1/2/4, IRF1, RELB) that are upregulated in response to IFNγ released by CD8+ T cells during immune response. Responders also showed more active cell-cell interactions. Overall, these observations suggest that responders had a more active immune response in the tumor even before treatment with immunotherapy. Interestingly, applying the model on on-treatment data showed significant increase of performance with respect to the pre-treatment point especially for models based on pathways and cell fractions (**Figure S4**). To investigate this improvement, we compared the distribution of the systems biomarkers pre- and on-treatment in responders and non-responders (**Figure 5H**). As expected, after treatment top positive biomarkers generally tended to increase in both responders and non-responders. This was the case, for example, for CD8^+^ T cells abundance, STAT1/2/4 and RFX-associated TFs, and IFNγ binding to the corresponding receptor. On the contrary, the fraction of ‘other’ (non-immune) cells decreases upon treatment, suggesting a reduction in tumor size. Overall, these observations are in agreement with an increased anti-tumor immunity upon treatment with immunotherapy.

### Integration of quantitative and spatial information on tumor infiltrating immune cells

Next, we performed preliminary analyses to evaluate whether the addition of information on spatial localization of Tumor-Infiltrating Lymphocytes (TILs) can improve the prediction of immune responses. We applied BEMKL and RMTLR on features related to local structural patterns of TILs computationally extracted from pathology images for 11 cancer types (Saltz et al., 2018). Using cross-validation on the TCGA data, we compared the predictive power of immune cell quantification alone and in combination with spatial features using both RMTLR and BEMKL algorithms (**Figure 6A**).

**Figure 6:**
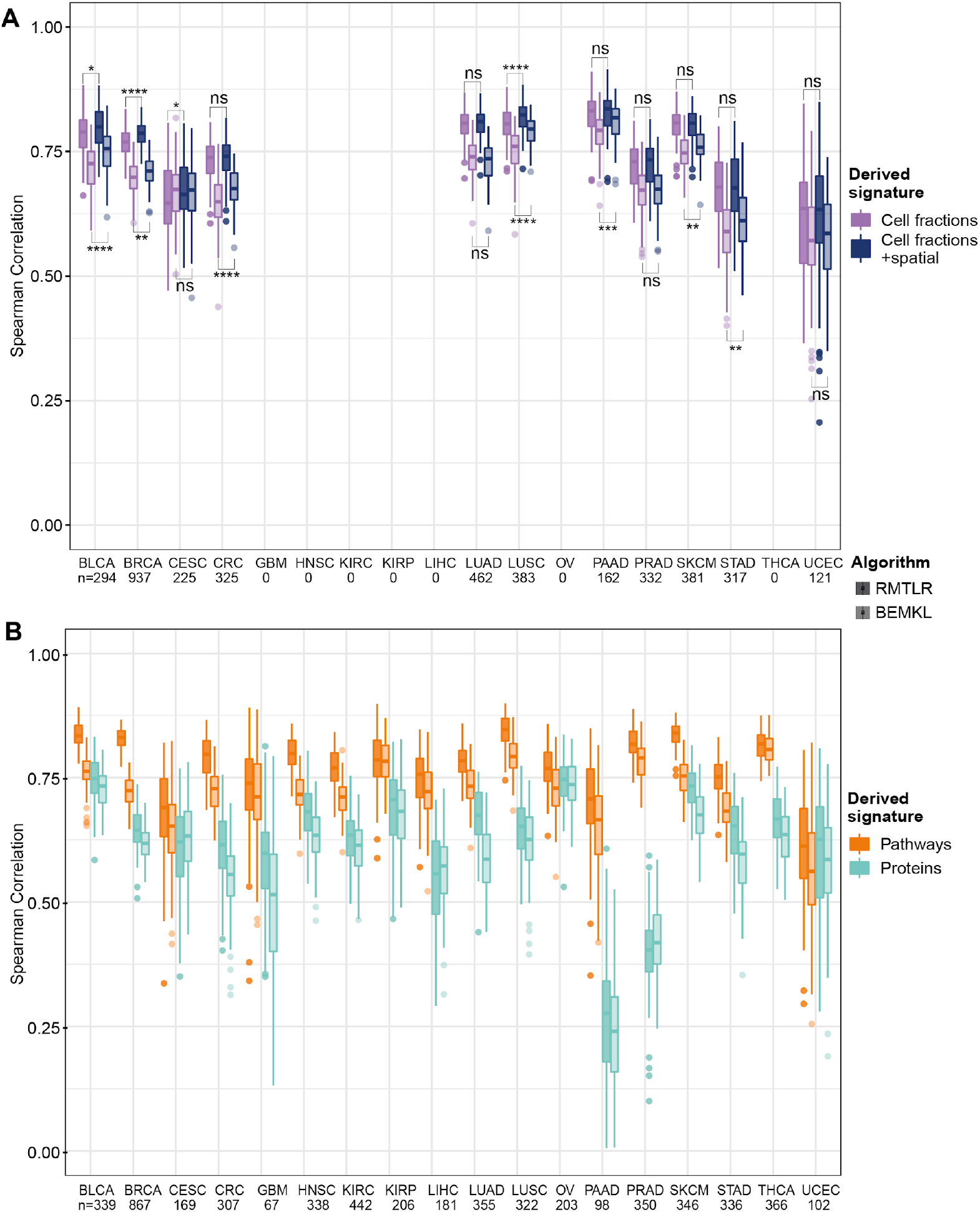
Comparison of predictive performance of different data modalities. RMTLR and BEMKL cross-validation prediction performances for (A) immune cell fractions alone, and in combination with spatial information of Tumor-Infiltrating Lymphocytes (TILs), and (B) protein expression and pathway activity. For each plot, we consider only cancer types and patients for which both data types were available (spatial information only available for 11 out of the 18 cancer types). Boxplots represent the distribution of prediction performances across the 100 randomized cross-validations. In (A) and (B), the prediction performance is evaluated using spearman correlation. The significance level (*p-value < 0.05, **p-value < 0.01, ***p-value < 0.001, ****p-value < 0.0001) indicates whether adding spatial information improves the performance with respect to using only immune cell fractions and whether pathway activity outperforms protein expression (one-side Wilcoxon signed-rank test).

The integration of information on the spatial localization of CD8^+^ T cells did not provide significant improvements in the performance for 4 and 7 of the 11 cancer types using RMTLR and BEMKL respectively (p-value > 0.05; **Figure 6A**). However, for bladder cancer (BLCA), breast cancer (BRCA), and lung (LUSC) squamous cell carcinoma, both algorithms showed a significant improvement (Wilcoxon sum-rank test, p-value < 0.05) in the prediction of the patient’s response when including the information of spatial localization of TILs (**Figure 6A**). As positive biomarkers of immune response we identified the percentage of TILs patches (in 8 out of 11 cancer types) and the Banfield-Raftery index (in 6 out of 11 cancer types), which is a measure related to the number of clusters (**Figure S5**). In fact, TILs need to be abundant and present in multiple clusters throughout the tissue in order to elicit an effective immune response (Galon and Bruni, 2019). These features were in general associated with better prognosis (Saltz et al., 2018). We identified the Ball Hall index, which is a measure of cluster dispersion, as a negative biomarker of immune response (in 6 out of 11 cancer types **Figure S5**). This is in agreement with the finding of Saltz et al, who associated higher Ball Hall index with less favourable survival (Saltz et al., 2018). While the automatic derivation of features describing the spatial localization of immune cells is still at its infancy, our results suggest their potential for prediction of response to ICB therapy.

### Pathway activity is a better predictor than proteomics

Using cross-validation of the TCGA data we further assessed the predictive power of proteomic profiling (expression of 200 proteins and 58 phosphoproteins) using reverse phase protein array (RPPA) data from The Cancer Proteome Atlas (TCPA) (Li et al., 2013). Cross-validation performances using proteomics data were compared with the corresponding performances of pathways activity (**Figure 6B**). The aim was to compare the predictive power of different types of quantification of intracellular pathways. Interestingly, pathway activity scores revealed higher ability to predict patients’ response to immunotherapy for all cancer types, except UCEC (p-value = 0.35), when compared with protein expression on the patients for which both data types were available.

## DISCUSSION

The efficacy of immunotherapy with immune checkpoint inhibitors depends on the intricate crosstalk across the cells in the TME. Thus, to disentangle the mechanisms underlying - and ultimately predicting- patients’ response, it is essential to adopt a holistic strategy to study patients’ tumors (Robert, 2020). To find effective biomarkers, it is essential to use a systems biology approach to investigate how different mechanisms contribute to the overall behavior, looking at the TME from different perspectives (Lapuente-Santana and Eduati, 2020; Senft et al., 2017). Towards this goal, in this study, we derived systems biomarkers of immune response considering the cellular composition of the TME together with inter- and intra-cellular communication to provide a more comprehensive and mechanistic characterization of tumors. Since RNA-seq data is becoming routinely available in clinical settings, we decided to focus on deriving system-based signatures by using prior knowledge to structure RNA-seq data into different mechanistic layers. Importantly, our approach proved to be effective in predicting responses to ICBs on independent datasets.

We used machine learning to look for associations between the derived system-based features and the immune response, estimated using 14 predictors (proxies) derived from recent publications. We considered these proxies as different tasks to be predicted by our machine learning models, and used multi-task learning algorithms in order to learn all tasks jointly. Multi-task techniques have the great advantage that they allow to exploit the shared information across tasks. By inducing regularization forcing the algorithms to perform well in all tasks, they prevent overfitting thereby providing more generalizable models. Another advantage of our approach is that it does not require a dataset where patients’ responses to immunotherapy are known for model training. These datasets are generally limited to a few dozen patients and are, therefore, not optimal for training of machine learning approaches due to risk of overfitting. Instead, we could exploit the large sample size of the TCGA RNA-seq dataset to build cancer-specific models, including cancer types for which the potential of ICB therapy has not been extensively studied yet. An additional advantage of our approach is that by deriving system-based signatures, it performs knowledge-guided dimensionality reduction (e.g. pathways consist of only 14 features derived from thousands of genes). Unlike other approaches for dimensionality reduction, which cause loss of interpretability of the derived features, our derived signatures actually improve the interpretability providing quantification of different complex mechanisms of the TME. By aggregating RNA-seq data into higher representations, our models provide views of the tumor that would have been accessible only through the generation of additional data with complex and expensive techniques (e.g. imaging for immune cell quantification, phosphoproteomics upon perturbation for pathway activation). Dimensionality reduction also allows improving algorithm performances reducing risk of overfitting.

In summary, our approach allows us to derive more generalizable models thanks to: 1. multi-task learning, leveraging info from multiple proxies of immune response; 2. training on large datasets from the TCGA; 3. dimensionality reduction using system-based features. This results in superior predictive power on completely independent datasets, outperforming also the gold standard computed as the average of the different literature-derived proxies of immune response.

An intriguing observation is that intracellular signaling pathways activity are major predictors of immune response. This is likely due to the fact that they regulate both tumor cell intrinsic (due to mutations) and extrinsic (due to exogenous stimulation) mechanisms of resistance to immune-attack. Importantly, in our analysis, we found that pathway activity was a better predictor than protein expression or phosphorylation. Limited predictive power of proteomics data might be partially due to the use of RPPA data, which is limited to a few hundreds of proteins and can be noisy. However, we believe that a main motivation for the superior performance of the pathway activity scores is that they were derived from perturbation-response signatures, therefore inferring the activity of the pathway by looking at how genes downstream of the pathways respond to perturbations (Schubert et al., 2018). This approach allows to take into account post-translational modification and capture the dynamic nature of the pathways even when only static RNA-seq data are available. Similar observations hold for TF activity, which was computed based on the expression of the regulated genes (Garcia-Alonso et al., 2019).

For both pathways and TFs, we identified biomarkers that could be used to suggest new therapeutic strategies. Intracellular networks regulate tumor cell interactions with the microenvironment via regulation of immune checkpoints, regulation of antigen presentation and release of inflammatory chemokines (Spranger and Gajewski, 2018). Targeting these intra-cellular networks in cancer cells has potential to improve efficacy of immunotherapy with ICB (Sharma and Allison, 2015) or as an alternative approach to immunotherapy by inhibiting the expression of immune checkpoints (Wu et al., 2019; Zerdes et al., 2018). For example, overactivation of the PI3K pathway, that we identified as a positive biomarker, was shown to cause overexpression of PD-L1 contributing to immune evasion (Coelho et al., 2017). PI3K inhibition resulted in downregulation of PD-L1 in different cancer types providing an alternative approach to ICB to enhance anti-tumor immunity (Zerdes et al., 2018). The VEGF pathway, identified instead as a negative biomarker of immune response in our analysis, has been already associated with immune exclusion and resistance to ICB therapy (Chen and Mellman, 2017). In line with our results, accumulating evidence suggests that combining ICB immunotherapy with antiangiogenic agents targeting VEGF might improve clinical efficacy of immunotherapy in patients with lung cancer (Manegold et al., 2017). Similarly, the TF PPARA, identified as a negative biomarker in our analysis, could be inhibited to augment inflammation in the TME. In line with this, a recent clinical trial showed that PPARA blockade promotes a more inflamed TME and improves ICB efficacy in advanced solid cancers (Laport et al., 2019). These results suggest that our biomarkers have strong potential to be exploited in future research to suggest new personalized therapies.

As expected, the tumor immune-cell composition was also important for predictions. However, complementing immune cell fractions with pathway information significantly improved the predictive power of both individual views. Furthermore, our preliminary analysis on TCGA data suggests that adding information on the spatial localization can improve the predictive performance in some cancer types. For this study we used data from one of the first attempts to automatically extract spatially relevant features for tumor tissue slides (Saltz et al., 2018). With the advent of clinical pathology, we expect spatial information to be increasingly available in clinical practice. As highlighted by our results, another important aspect for prediction of immune response is the intercellular communication. Although these types of communications are still less explored, our results suggest that they deserve more attention in future research. In this regard, more refined approaches to infer cell-cell networks from bulk RNA-seq data, and the exploitation of orthogonal information from single-cell technologies will be extremely valuable (Armingol et al., 2021).

Remarkably, we found literature support validation for most of our top biomarkers. This highlights the potential of using a systematic and unbiased approach like the one described in this paper. Instead of focusing on individual mechanisms requiring specific biological assays, we use widely available RNA-seq data complemented by prior knowledge to provide a holistic picture of the TME. In this way, we provide a tool (EaSIeR) that can be readily used to predict individual patients’ response to ICB therapy and paves the way to suggest new therapeutic strategies not only for individual tumor types but also for individual patients, based on systems biomarkers.

## METHODS

### TCGA RNA sequencing data

Gene expression data for 18 solid tumors were downloaded via the Firehose tool from the BROAD Institute (https://gdac.broadinstitute.org), released January 28, 2016. We selected primary tumor or metastatic (only in the case of melanoma) samples, resulting in a total of 7750 patients.

We extracted the gene expression data from “illuminahiseq_rnaseqv2-RSEM_genes” files. From this data, we used “raw_count” values as counts and we calculated Transcripts Per Millions (TPM) from “scaled_estimate” values multiplied by 1M. We first removed those genes with a non-valid HGNC symbol and we averaged the expression of those genes with identical HGNC symbols.

### Validation data

Validation cohorts for melanoma and gastric carcinoma were derived from published datasets including both response to ICB therapy and RNA sequencing (RNA-seq) raw data (Auslander et al., 2018; Gide et al., 2019; Kim et al., 2018). We considered only patients treated with anti-PDL1. On the basis of Response Evaluation Criteria in Solid Tumors (RECIST), we considered responders (R) patients with complete response (CR) or partial response (PR), and non-responders (NR) patients with progressive disease (PD) or stable disease (SD). More details can be found in **Table S5**. For the three datasets, we downloaded the corresponding SRA files from the Sequence Read Archive (SRA, https://www.ncbi.nlm.nih.gov/sra/) and converted to FASTQ using the “fastq-dump function” provided by the SRA toolkit. FASTQ files of RNA-seq reads were then pre-processed with quanTIseq to obtain gene counts, TPM, and cell fractions (Finotello et al., 2019b). In brief, we used Trimmomatic (Bolger et al., 2014) to remove adapter sequences and read ends with Phred quality scores lower than 20, discard reads shorter than 36 bp, and trim long reads to a maximum length of 50 bp (quanTIseq preprocessing module). We run Kallisto (Bray et al., 2016) on the pre-processed RNA-seq reads to generate gene counts and transcripts per millions (TPM) using the “hg19_M_rCRS” human reference (quanTIseq gene-expression quantification module).

### System-based signatures of the TME

We used RNA-seq data to derive different types of mechanistic signatures integrating prior knowledge.

#### Immune cell quantification

We used quanTIseq to compute tumor-infiltrating immune cell fractions which are estimated by applying deconvolution to bulk gene expression levels in a mixture based on cell-specific gene expression signatures (Finotello et al., 2019b). quanTIseq returns the fractions of 10 cell types: B cells, classically (M1) and alternatively (M2) activated macrophages, monocytes, neutrophils, natural killer cells, non-regulatory CD4^+^ T cells, CD8^+^ T cells, regulatory T (T_reg_) cells and myeloid dendritic cells. The fraction of other cell types in the mixture is computed as one minus the total fraction of immune cells and was shown to often correlate with tumor purity (Finotello et al., 2019b). Since non-regulatory CD4^+^ T cells are difficult to distinguish from T_reg_ cells, we decided to consider non-regulatory CD4^+^ T cells as the sum of both non-regulatory CD4^+^ T cells and T_reg_ cells, keeping T_reg_ cells as a separated cell type as well.

#### Pathway activity

We used PROGENy to compute scores for 14 pathways: Androgen, EGFR, Estrogen, Hypoxia, JAK-STAT, MAPK, NFkB, p53, PI3K, TGFb, TNFa, Trail, VEGF and WNT (Holland et al., 2020; Schubert et al., 2018). Pathway-specific signatures were derived from pathway perturbation experiments by investigating which genes change in expression when a pathway is perturbed. A linear regression model is used to fit the genes that are affected by the perturbation of the pathway. Both pathway-specific signatures and gene expression data are then used to infer pathway signaling activity. The pathway scores were directly computed using the progeny R package version 1.10.0. Since these scores are actually a linear transformation of gene expression data, we therefore removed 448 genes used to compute the proxies of immune response (average pan-cancer Pearson correlation with original pathway activity = 0.99, p-value < 10^−16^).

#### Transcription factor activity

We used DoRothEA to compute TF activity (Garcia-Alonso et al., 2019). The expression of individual genes is controlled by TFs, and TF activity can be estimated by the expression of its target genes (so-called TF regulons). The TF activity was estimated using analytic Rank-based Enrichment Analysis (aREA) from the viper R package 1.22.0, as part of the dorothea R package 1.0.1. aREA provides a normalized enrichment score for each TF regulon based on the average ranks of its targets. Each TF-target interaction is assigned a degree of confidence (from A to E) depending on the total of supporting evidence. To only consider high quality regulons, we filtered for confidence levels A and B resulting in a total of 115 TFs.

#### Ligand-receptor pairs

Based on ligand-receptor (LR) pair annotations from the database by Ramilowski et. al. (Ramilowski et al., 2015) we quantified LR interactions in the TME for each individual patient. This was done in two steps: first we derived a subset of 867 LR pairs that are potentially present in the TME, then we quantified these pairs for each patient based on RNA-seq data.

For the first step we started from the 1894 LR literature supported pairs in the Ramilowski database, consisting of 642 unique ligands and their 589 cognate receptors. Furthermore, the database annotates the TPM expression of these ligands and receptors in 144 human cell types based on CAP Analysis of Gene Expression (CAGE) from the FANTOM5 expression atlas. We filtered for the 24 cell types commonly acknowledged to be present in the TME and present in the Ramilowski database (**Table S6**). Additionally, we considered a pan-cancer cell type derived using data from the cancer cell line encyclopedia (CCLE) (Barretina et al., 2012). In CCLE we selected gene expression data for all cell lines linked to the 18 solid cancer types researched here, leaving 583 cell lines. We determined the median expression of each gene over all selected cell lines, which we considered as the gene expression of the pan-cancer cell type. We filtered for ligands and receptors with expression ≥ 10 TPM in at least one of the 25 cell types considered. Furthermore, we excluded ligands and receptors that were expressed by a cell type but not paired to another ligand or receptor in one of the other 24 considered cell types, resulting in 867 LR pairs. The 10 TPM threshold used in the Ramilowski paper for the CAGE data was based on known expression data from B-cells. To confirm the suitability of the 10-TPM threshold for the CCLE RNA-seq data, we considered six healthy B-cell datasets from two studies (Hoek et al., 2015; Linsley et al., 2014). By comparing the sets of ligand-receptor pairs expressed in the Ramilowski’s CAGE data versus RNA-seq B-cell considering different thresholds. The 10-TPM cut-off allowed the retrieval of ~80% B cell-specific ligand-receptors expressed in the Ramilowski’s data, while resulting in the RNA-seq-specific expression of only 3% of the full ligand-receptors set (data not shown).

Next, we assigned a patient-specific weight to the LR pairs. The LR pair weight was defined as the minimum of the log2(TPM+1) expression of the ligand and the receptor, theorizing that a pair has a weaker bond if one of the genes is expressed at a lower level.

#### Cell-cell interactions

The 24 TME cell types were combined in 12 aggregated cell types (**Table S6**). To assign a weight to the cell-cell (CC) interactions we considered the number of active LR pairs between each pair of the 13 considered cell types (12 TME aggregated cell types and the additional pan-cancer cell type), for a total of 169 cell-cell pairs. For each CC pair (sender→receiver cell) we considered a LR pair active only if the ligand is expressed in the sender cell and the receptor is expressed in the receiver cell, using the 10 TPM threshold as described above. For each LR pair we computed the frequency across the whole TCGA database, with the idea that more rare interactions are more relevant to discriminate patients. The cell-cell score for each patient is then computed as the sum of the inverse of the frequency of all the active ligand-receptor pairs.

### Spatial localization of Tumor-Infiltrating Lymphocytes (TILs) extracted from imaging

Images of tumor-tissue slides are also available for some of the TCGA patients. We used the spatial information of TILs (Saltz et al., 2018) derived from applying deep learning on digitized H&E-stained images. For model training we used 6 out of the 10 spatial features provided in the original publication, thus excluding highly correlated features (**Figure S6**). The features considered are: TIL percentage, within-cluster dispersion (WCD) mean, Ball-Hall adjusted, Banfield-Raftery adjusted, C index adjusted, and determinant ratio adjusted. TIL percentage correlates with the number of positive TIL patches, the number of TIL clusters and the cluster size mean. The Ball-Hall index correlates with the cluster extent mean. The Banfield-Raftery index correlates with the number of TIL clusters.

### Proteomics data

We used protein data for 18 solid tumors for a total of 5394 patients. Data were downloaded via The Cancer Proteome Atlas Portal (https://tcpaportal.org). We used reverse phase protein array (RPPA) data labeled “Level 4 Pan-Can 32”, including 200 proteins and 58 phosphoproteins for a total of 258 features. For RMTRL, protein features with any missing value in a specific cancer type were not considered (in the range from 24 to 47 proteins depending on the cancer type). This was not done for BEMKL, which can handle missing values.

### Transcriptomics based scores of immune response

The 14 published transcriptomics signatures of immune response are summarized in **Table S1**. Among them, immune cytolytic activity (Rooney et al., 2015) represents the level of two cytolytic effectors granzyme A (GZMA) and perforin (PRF1) which are overexpressed upon CD8+ T cell activation, Ock immune signature (Ock et al., 2017) is based on the expression of 105 genes associated with response to immunotherapy with MAGE-A3 antigen. Immunophenoscore (Charoentong et al., 2017) is calculated according to genes related to MHC molecules, immunomodulators, effector and suppressor cells. IMPRES (Auslander et al., 2018) is obtained through a logical comparison between the expression of immune checkpoint gene pairs. Roh immune score (Roh et al., 2017) is defined by a set of genes involved in immune activation in relation to tumor rejection. Chemokine signature (Messina et al., 2012) is based on a gene set associated with inflammation and immunity, that is able to predict host immune reaction and the formation of tumor-localized lymphoid structure. Davoli immune signature (Davoli et al., 2017) is derived from the expression of cytotoxic CD8+T cells and NK cells markers. IFNγ signature (Ayers et al., 2017) comprises genes being able to separate responders and non-responders in melanoma Immune expanded signature (Ayers et al., 2017) is generated searching for genes highly correlated with IFNγ signature genes, this new set included all immune-related genes T-cell inflamed microenvironment signature (Ayers et al., 2017) is based on the joint potential of IFNγ and T cell-associated inflammatory genes in predicting response to PD-1 blockade. TIDE (Jiang et al., 2018) is developed on the basis of immune escape signatures such as T cell dysfunction or exclusion. MSI status (Fu et al., 2019) is determined by logical comparison of MSI-related gene pairs.

All these proxies were calculated following the methodology reported by original studies. Only for the chemokine signature, we adjusted the sign according to the positive correlation with the cluster of correlated tasks. Since it is computed based on the first principal component, the sign is arbitrarily determined. When applicable, these transcriptional predictors were computed according to published computational frameworks. More details can be found in (**Table S1**).

### Machine learning methods

#### Regularised Multi-Task Linear Regression (RMTLR)

The objective function that defines the RMTLR for observations is describe in Equation 1:

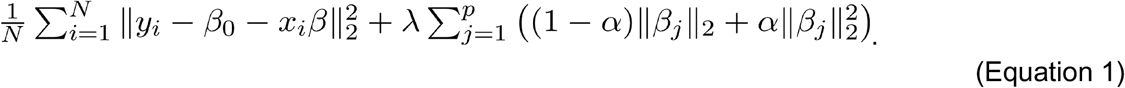

In this equation, *y*_*i*_ represents a *q*-dimensional row-vector where each entry corresponds to a task, and is a row vector where each entry represents an observed feature. The aim of the RMTLR is to estimate a matrix *β* whose rows represent the relation between one feature and all the tasks, and a vector *β*_0_ of offsets (one for each task). The regularization term of RMTLR is a grouped version of the elastic net that aims at enforcing sparsity to entire rows of *β* (Yuan and Lin, 2006b; Zou and Hastie, 2005). In this way, the features corresponding to those rows of *β* that are set to zero do not contribute to the model. The strength of the regularization effect is tuned via the hyperparameter *λ*, while *α* regulates the interplay between the ridge- and the lasso-like terms of the elastic net. We selected the hyperparameters using 5-fold cross-validation.

We used RMTLR as implemented in the glmnet R package 2.0-16 (Friedman et al., 2010). When applying RMTLR to a combination of multiple views, individual derived signatures (single views) were combined by merging datasets by columns.

#### Bayesian Efficient Multiple Kernel Learning (BEMKL)

BEMKL (Gönen, 2012) is a Bayesian approach with two important features: multi-view and multi-task learning. BEMKL is a non-linear regression model which defines view-specific kernels as similarity measures between all samples and integrates them into a combined kernel to obtain response predictions (Equation 2). The similarity between samples is calculated using the gaussian kernel.

On the one hand, BEMKL uses multi-view learning to integrate different sample views as kernels, creating a combined kernel as the weighted sum of the view-specific kernels. The kernel weights were learned using multiple kernel learning and represent the view’s importance for predicting the response. On the other hand, the peculiarity of multi-task learning is that it enables to model multiple tasks simultaneously. Assuming that the kernel weights are shared across all tasks, task-specific weights are estimated for all samples.

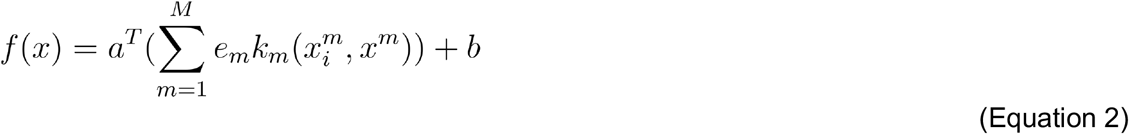

Where M denotes the number of input kernels, 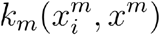 represents the view-specific kernel, *e*_*m*_ the shared kernel weights *α* and the task-specific sample weights.

Bayesian inference was used to estimate all model parameters that were interpreted as random variables with certain probability distributions. The parameters of these distributions were learned using deterministic variational approximation. More detailed explanation of the probabilistic model used, and the inference method can be found in the original paper (Costello et al., 2014; Gönen, 2012). An R implementation of this method was available at https://github.com/mehmetgonen/bemkl. An adaptation to multi-task learning was available at https://github.com/mehr-een/bemkl-rbps and we converted the code from Matlab to R https://github.com/olapuentesantana/easier_manuscript.

#### Model training based on TCGA data

Models were trained using the TCGA data, separately for each cancer type. For both RMTLR and BEMKL training was repeated 100 times with randomized cross-validation, each time randomly picking 20% of the samples as a test set. This helped to assess the stability of the model, in terms of both performance and feature selection. For each iteration, we first standardized the training set, and then we standardized the test set based on the mean and standard deviation of the training set.

#### Definition of biomarkers of immune response

As mentioned above, the model training was repeated 100 times which allowed us to assess the stability of the features. As a result, we estimated 100 weight values for each biomarker. The values displayed in Figures 2, 3 and 4 were defined as the median of the estimated weights, first across runs and second across tasks (median > 0, i.e. selected by regularization in at least 50% of the runs).

#### Prediction of response to ICB therapy

All the 100 models learned in the randomized cross-validation were included in the EaSIeR tool and were used to make predictions for the external test tests. Predictions for each task were computed as the average of the 100 models.

### Statistical analysis

We used one-side Wilcoxon signed-rank test (pair data) for comparison of predictions between pairwise combinations of derived signatures and single ones; and two-sided Wilcoxon-rank sum test (unpaired data) for biomarker comparison between responders and non-responders. Statistical tests were carried out using the function ‘wilcox_test’ from the R package rstatix version 0.6.0. Effect size was calculated as the test statistic divided by the square root of the number of observations, using the function ‘wilcox_effsize’ from the R package rstatix version 0.6.0.

Model performances were evaluated using Spearman correlation for randomized cross-validation. For the validation dataset we computed Receiver-Operating-Characteristic (ROC) curve and Area Under the Curve (AUC) using the R package ROCR 1.0-11, were used.

All analyses were performed using R software, version 4.0.2. For training of machine learning models, we used R software version 3.5.2.

## Data and code availability

All the datasets used are publicly available (**Table S5**). The code used for model training and analysis is available at https://github.com/olapuentesantana/easier_manuscript, and the EaSIeR to compute system biomarkers and likelihood of patient response to ICB from RNA-seq data is available at https://github.com/olapuentesantana/mechanistic_biomarkers_immuno-oncology.

## ACKNOWLEDGEMENT

We thank Federico Marini for useful discussion and feedback. F. F. was supported by the Austrian Science Fund (FWF) (project No. T 974-B30).

## AUTHOR CONTRIBUTION

Conceptualization F.E. and F.F.; Methodology and Formal analysis O.L.S., M.G., F.F. and F.E.; Software O.L.S.; Results interpretation O.L.S, F.E., F.F., P.H; Writing - Original Draft F.E., O.L.S. and F.F.; Review & Editing O.L.S, M.G., P.H., F.F., F.E.

