## Supplementary Information for "Predictive systems biomarkers of response to immune checkpoint inhibitors"

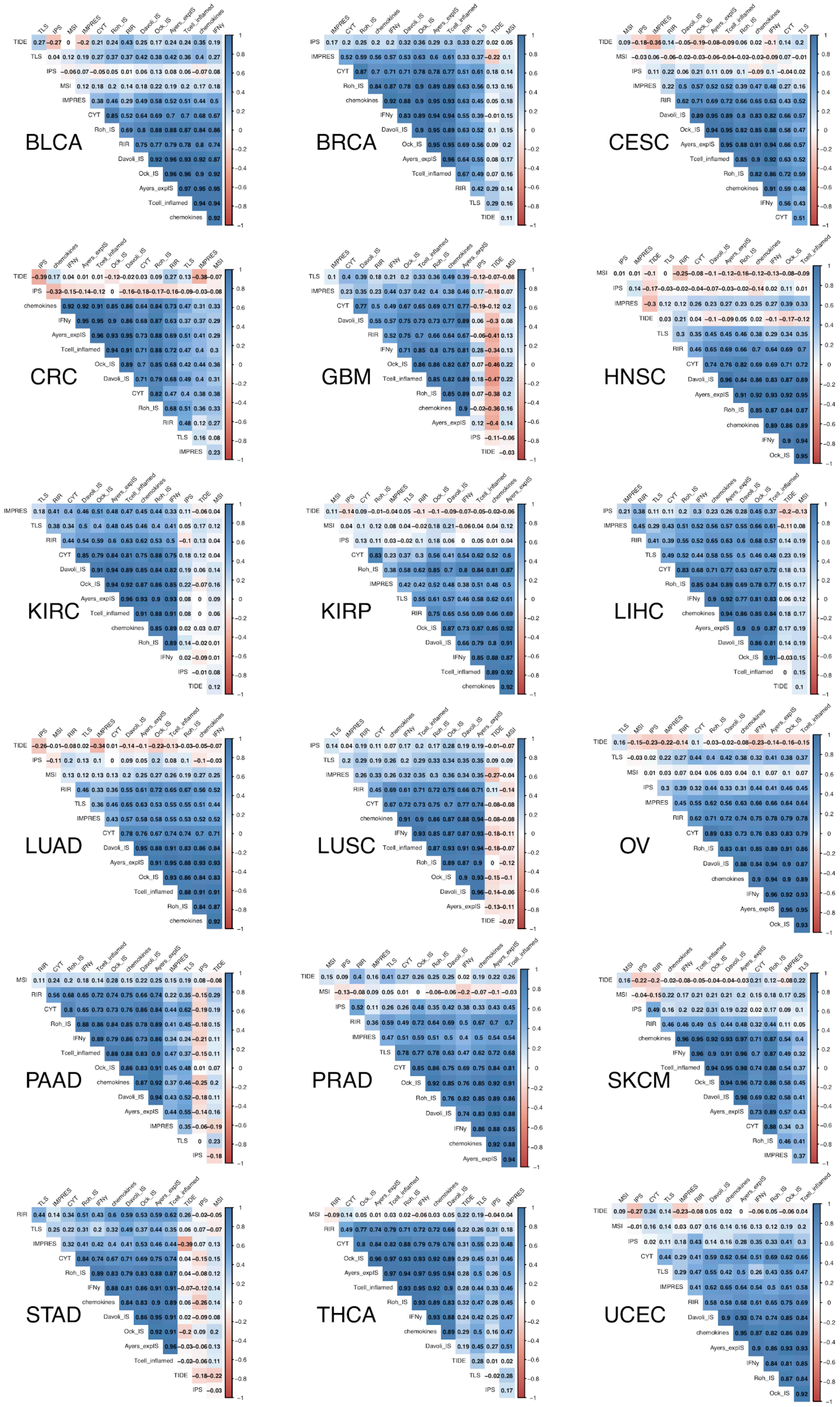

Figure S1. Cancer-specific Pearson correlations between the 14 proxies of immune response.

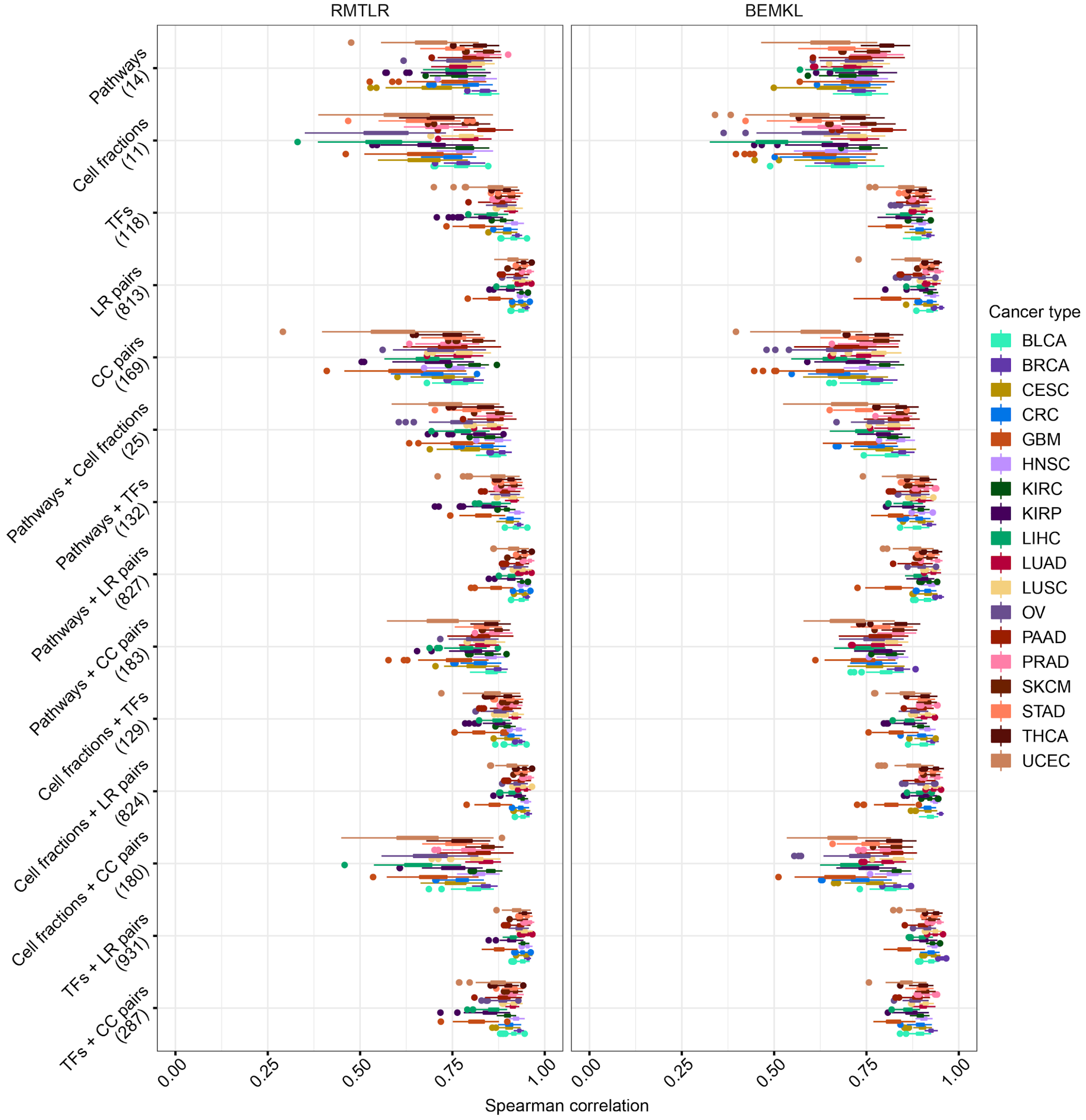

Figure S2. Randomized cross-validation performance of cancer-specific models trained on the TCGA patients. Models performances are shown for models trained separately for each system-based signature (single view) and for pairwise combinations of views. For each input data, the number of features is given in brackets.

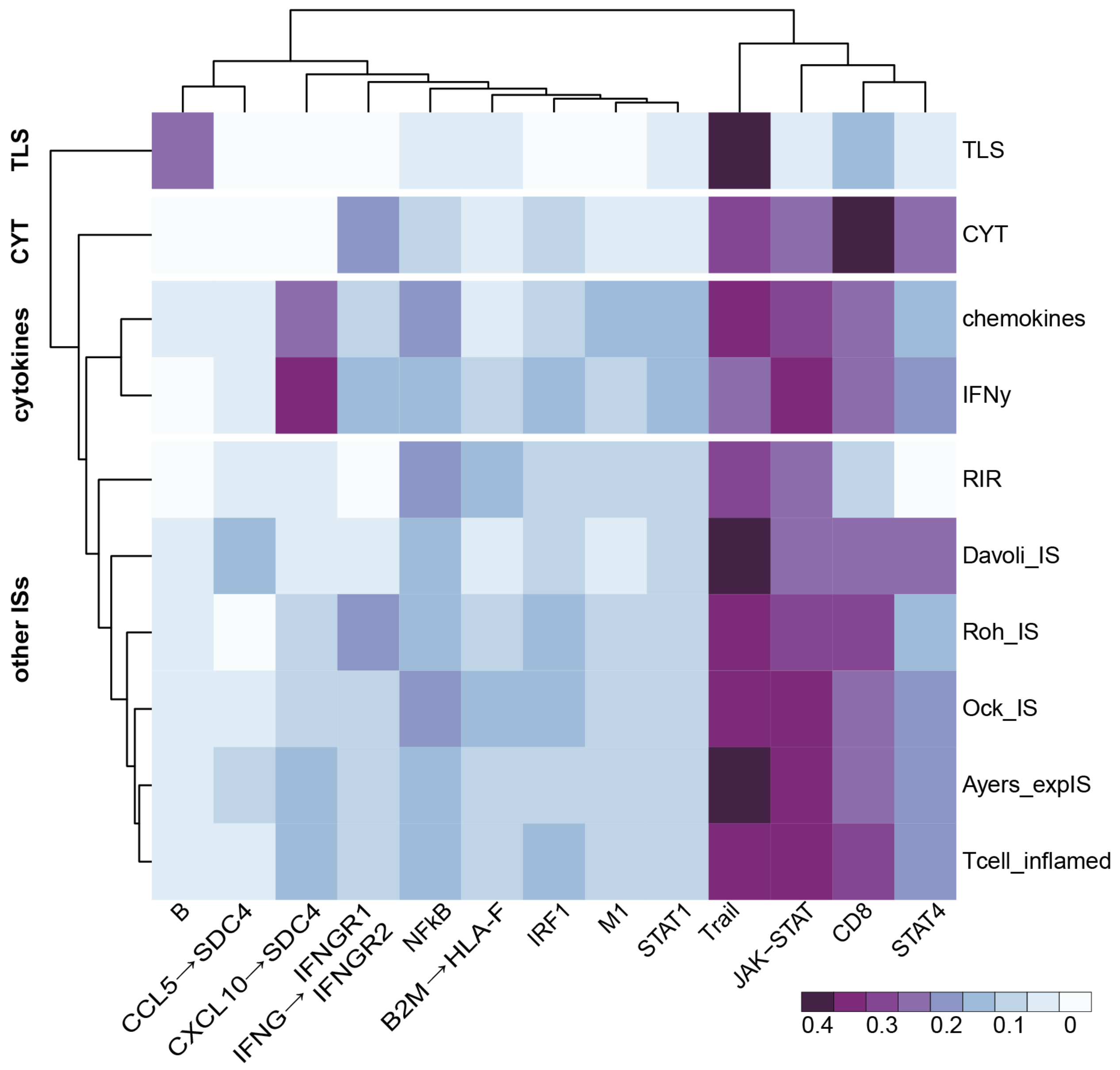

Figure S3. Heatmap showing the median across cancer types of the estimated weight of the features for each task. Only features with variance across tasks ≥ 0.0015 (top 1%) are shown in the heatmap. Tasks cluster in four main groups: 1. Tertiary lymphoid structures (TLS), 2. cytolytic activity (CYT), 3. cytokines related proxies (chemokines and IFNγ) , 4. all other immune signatures (ISs).

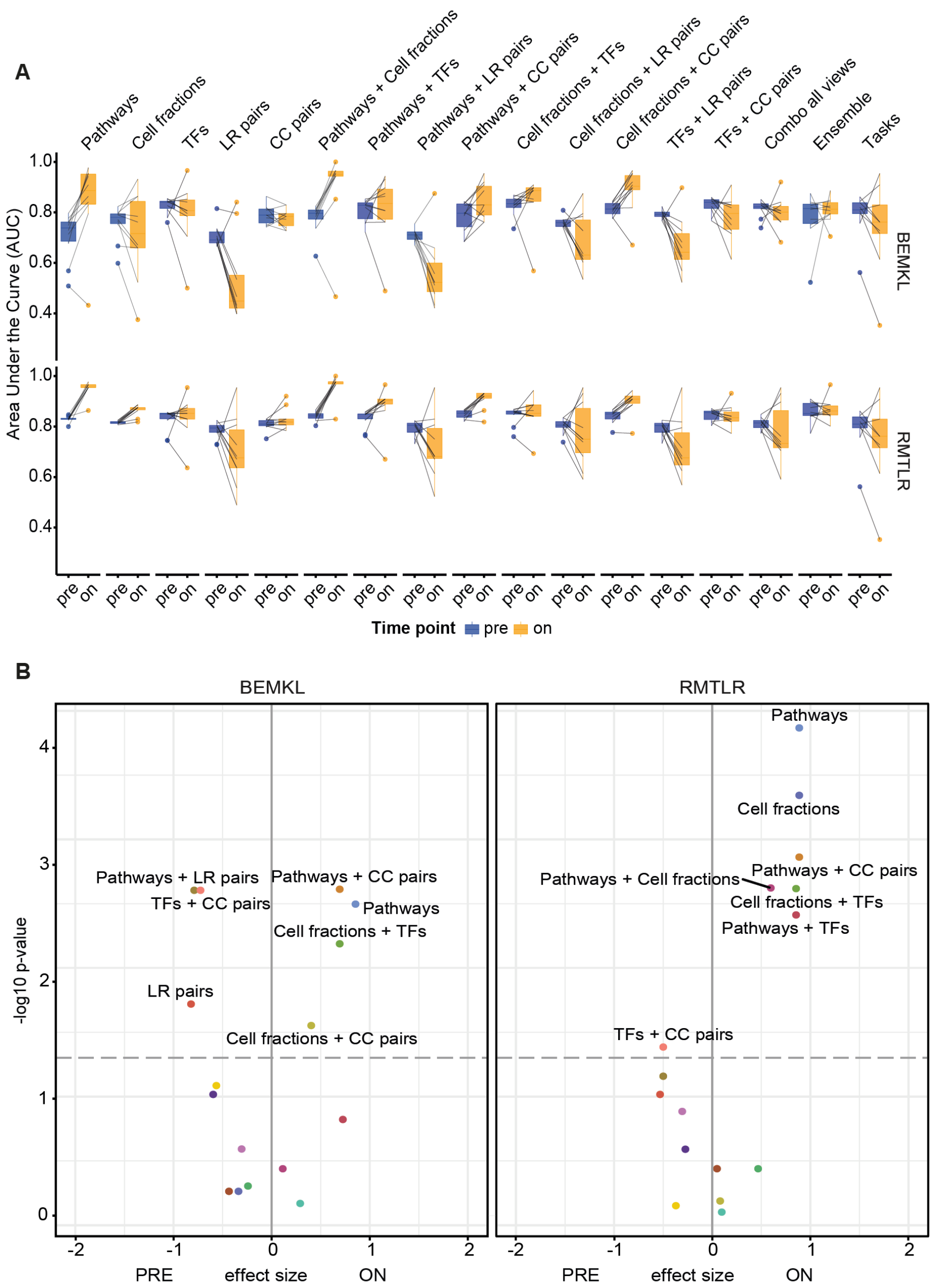

Figure S4. (A) Comparison of Area Under the Curve (AUC) values for the different tasks computed on the pre-treatment (PRE) and on-treatment (ON) samples for the melanoma dataset (Auslander et al., 2018; Gide et al., 2019). Performances are shown for single views, pairwise combinations of views, combination of all views, average of single views predictions (ensemble), and all tasks (gold standard). (B) Volcano plots showing the statistical comparison of pre- vs on-treatment samples (two-sided Wilcoxon-rank sum test). Results are shown for both BEMKL and RMTLR algorithms.

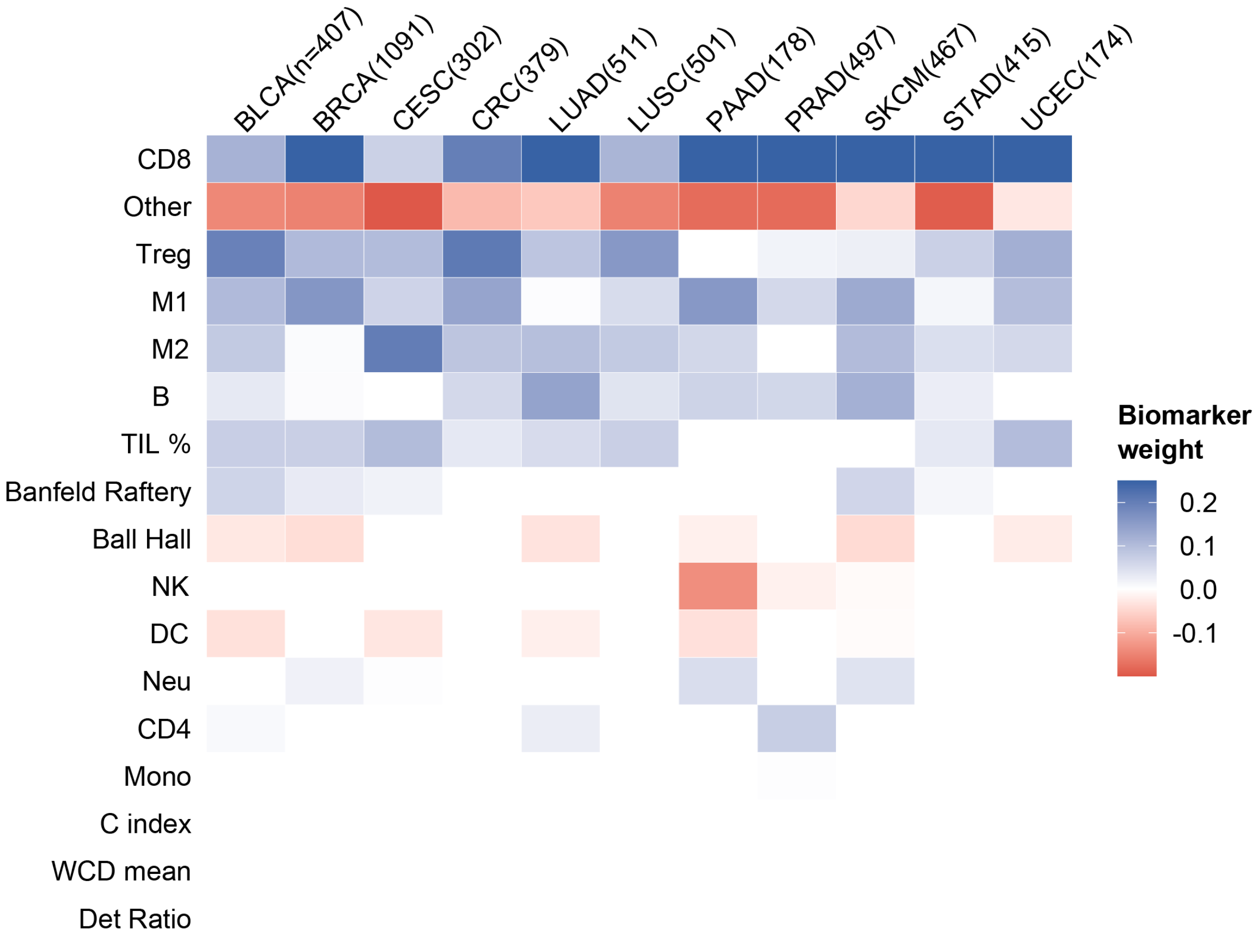

Figure S5. Heatmap showing regression coefficients for cancer type-specific models when using immune cell fractions in combination with spatial information of TILs. Shown are the median values computed first across 100 randomized cross-validation runs (to keep only robust biomarkers) and then across tasks. Positive (blue), none (white) or negative (red) relationship of each biomarker with the immune response. Rows (biomarkers) were sorted according to their absolute mean value across tumors.

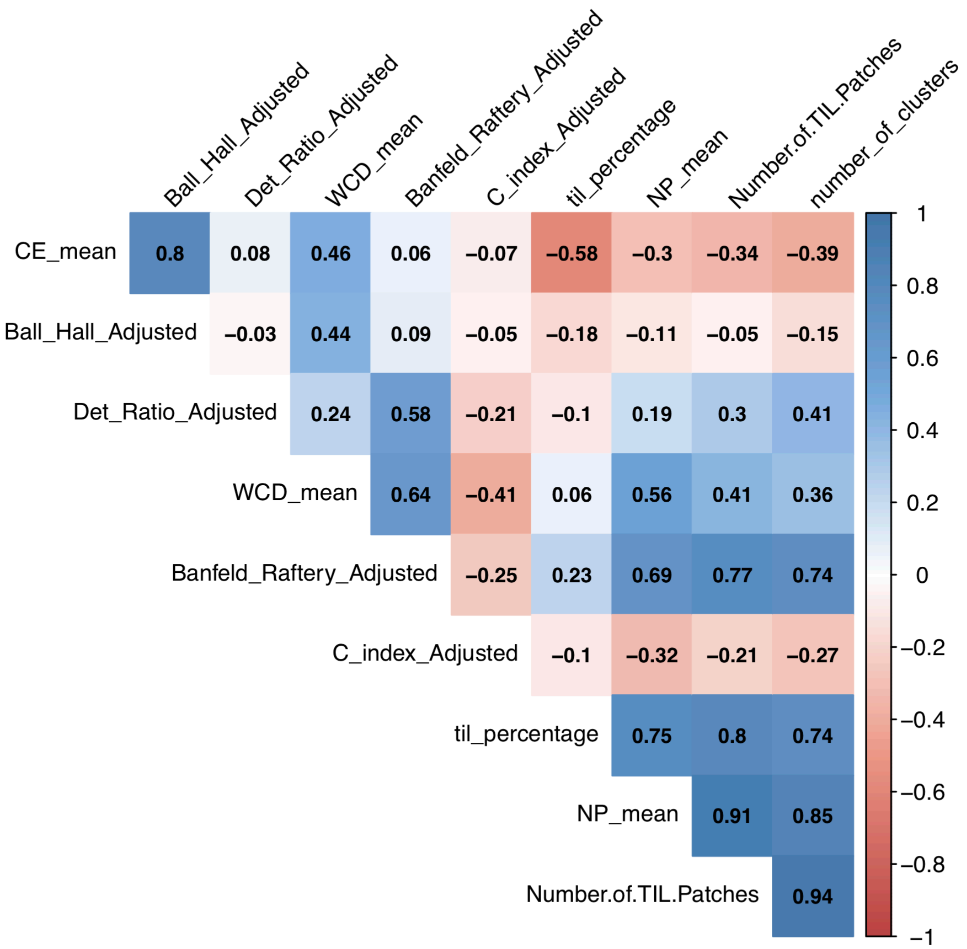

Figure S6. Pan-cancer correlation of spatial features described in (Saltz et al., 2018).

Table S5. Immunotherapy published studies (with available bulk RNA-Seq) information.

| **Original study** | **Cancer type** | **Prior therapies** | **Biopsy** | **Samples used** | **R and NR** | **RNA-seq fastq files** |
|---|---|---|---|---|---|---|
| Auslander (Auslander et al., 2018) | Melanoma (Metastasis) | Therapy naive | Fresh-frozen | PD-1:<br>- Pre: n=9 (R=1, NR=8)<br>- On: n=17 (R=0, NR=17) | As reported in publication. | PRJNA476140 |
| Gide (Gide et al., 2019) | Melanoma (Metastasis) | BRAFi | FFPE | PD-1:<br>- Pre: n=41 (CR=4, PR=15, PD=16, SD=6)<br>- On: n=9 (CR=0, PR=4, PD=4, SD=1) | R= CR, PR<br>NR= SD, PD | PRJEB23709 |
| Kim (Kim et al., 2018) | Gastric cancer (Metastasis) | Prior failure of at least 1 line of chemotherapy (platinum) | Fresh-frozen | Pre: n=45 (CR=3, PR=9, PD=18, SD=15) | R= CR, PR<br>NR= SD, PD | PRJEB25780 |

FFPE: Formalin-fixed paraffin-embedded
CR: Complete Responder; PR: Partial Responder; PD: Progressive Disease; SD: Stable Disease
R: Responder; NR:Non-responder

Table S6. List of cells included in the TME network.

| Aggregate cell type | Cells in the literature | Cell types in network | Derivation |
|---|---|---|---|
| **Adipocytes** | Adipocytes (Balkwill et al., 2012; Wang et al., 2017) | Mature Adipocyte<br>Adipocyte Omental | |
| **B-cells** | B-cell (Fridman et al., 2012)<br>CD20 B-cell (Fridman et al., 2017; Wang et al., 2017)<br>CD19 B-cell (Wang et al., 2017) | CD19+ B cells | |
| **CD4+ T cells** | CD4+ T helper cells (Balkwill et al., 2012; Fridman et al., 2012; Wang et al., 2017)<br>CD4+ T regulatory cells (Balkwill et al., 2012; Fridman et al., 2012) | CD4+ T cells<br>CD4+CD25+CD45RA+ naive regulatory T-cells<br>CD4+CD25+CD45RA- memory regulatory T-cells<br>CD4+CD25-CD45RA+ naive conventional T-cells<br>CD4+CD25-CD45RA+ memory conventional T-cells | CD4+ NR T cells<br>CD4+ MR T cells<br>CD4+ NC T cells<br>CD4+ MC T cells |
| **CD8+ T cells** | CD8+ cytotoxic T cells (Balkwill et al., 2012; Fridman et al., 2012, 2017; Wang et al., 2017) | CD8+ T cells | |
| **Dendritic cells** | Dendritic cell (Balkwill et al., 2012; Fridman et al., 2012, 2017; Wang et al., 2017) | Dendritic Plasmacytoid<br>Dendritic Monocyte Immature derived | Dendritic Monocyte Id |
| **Endothelial cells** | Endothelial cells (Fridman et al., 2012, 2017)<br>Vascular endothelial cells (Balkwill et al., 2012)<br>Lymphatic endothelial cells (Balkwill et al., 2012) | Endothelial Microvascular<br>Endothelial Lymphatic | |
| **Fibroblasts** | Fibroblasts (Balkwill et al., 2012; Fridman et al., 2012, 2017; Wang et al., 2017) | Fibroblast Lymphatic<br>Fibroblast Skin Normal | |
| **Macrophages** | M1 macrophage (Balkwill et al., 2012; Fridman et | Macrophage Monocyte derived | Macrophage Monocyte |

| | | |
|---|---|---|
| | al., 2012, 2017; Richards et al., 2013; Wang et al., 2017)<br>M2 macrophage (Balkwill et al., 2012; Fridman et al., 2012; Richards et al., 2013; Wang et al., 2017) | |
| **Mast Cells** | Mast cells (Fridman et al., 2012, 2017; Wang et al., 2017) | Mast cells<br>Mast cells stimulated |
| **Monocytes Myeloid** | Myeloid derived suppressor cells (Fridman et al., 2012, 2017)<br>Monocytes (Fridman et al., 2017; Richards et al., 2013; Wang et al., 2017) | CD14+ Monocytes<br>CD14+CD16+ Monocytes<br>CD14+CD16- Monocytes<br>CD14-CD16+ Monocytes |
| **NK Cells** | Natural killer cells (Balkwill et al., 2012; Fridman et al., 2012, 2017; Wang et al., 2017)<br>Natural killer T-cells (Balkwill et al., 2012) | NK cells |
| **Neutrophils** | Neutrophils (Balkwill et al., 2012; Fridman et al., 2017) | Neutrophils |
